# The future impact of climate and land-use changes on Anatolian ground squirrels under different scenarios

**DOI:** 10.1101/2021.09.14.460244

**Authors:** Hakan Gür

**Affiliations:** Anatolian Biogeography Research Laboratory, Kırşehir Ahi Evran University, Kırşehir, Turkey

**Author notes:** Corresponding author. *E-mail address:* (H. Gür).

**Keywords:** Anatolia, biodiversity hotspots, biodiversity loss, climate change refugia

## Abstract

Climate and land-use changes are among the most important drivers of biodiversity loss and, moreover, their impacts on biodiversity are expected to increase further in the 21st century. In this study, the future impact of climate and land-use changes on Anatolian ground squirrels was assessed. Accordingly, a hierarchical approach with two steps was used. First, ecological niche modelling was used to assess the impact of climate change in areas accessible to Anatolian ground squirrels through dispersal (i.e. the impact of climate change). Second, based on the habitat preferences of ground squirrels, land-use data were used to assess the impact of land-use change in suitable bioclimatic areas for Anatolian ground squirrels under present and future conditions (i.e. the combined impact of both changes). Also, priority areas for the conservation of Anatolian ground squirrels were identified based on *in-situ* climate change refugia. This study represents a first attempt to combine niche modelling and land-use data for a species in Anatolia, one of the most vulnerable regions to the drivers of biodiversity loss, because it is the region where three of biodiversity hotspots meet, and interact. Habitat availability (i.e. suitable habitats across suitable bioclimatic areas) was projected to decline by 19-69% in the future (depending on the scenario), mainly due to the loss of suitable bioclimatic areas (47-77%, depending on the scenario) at lower elevations and in the western part of the central Anatolia and in the eastern Anatolia, suggesting that Anatolian ground squirrels will contract their range in the future, mainly due to climate change. Thus, *in-situ* climate change refugia were projected mainly in the eastern and southeastern parts of the central Anatolia, suggesting these regions as priority areas for the conservation of Anatolian ground squirrels.

## 1. Introduction

The rate of global change in nature during the past 50 years is unprecedented in the history of human life. The five greatest drivers of global change in nature and therefore ongoing biodiversity loss (e.g. around one million species already face the risk of extinction) are: changes in land and sea use, direct exploitation of organisms, climate change, pollution, and invasion of alien species (IPBES 2019). That is, climate and land-use changes are among the most important drivers of biodiversity loss and, moreover, their impacts on biodiversity are expected to increase further in the 21st century. For example, by the year 2100, at 1.5°C of global warming (above pre-industrial levels), 4% (2-9%) of 12,640 species of amphibians, reptiles, birds, and mammals are projected to lose over half of their climatically determined ranges, and at 4.5°C, this increases to 44% (31-59%) (Hoegh-Guldberg et al. 2018, Warren et al. 2018). Moreover, by the year 2070, substantial declines in suitable habitats are identified for approximately 19,400 species of amphibians, birds, and mammals, with approximately 1,700 species expected to become imperiled due to increased human land-use (Powers and Jetz 2019).

A region should meet two main criteria to be identified as a biodiversity hotspot. A biodiversity hotspot should have (1) ≥ 1,500 endemic vascular plant species and (2) ≤ 30% of its original natural vegetation. In other words, it should be irreplaceable and threatened. According to these criteria, 36 biodiversity hotspots (i.e. regions “where exceptional concentrations of endemic species are undergoing exceptional loss of habitat”, Myers et al. 2000) were identified around the world. The forests and other remnant habitats in these biodiversity hotspots correspond to only 2.5% of Earth’s land surface, but still have more than 50% of endemic plant species and nearly 43% of endemic amphibian, reptile, bird, and mammal species in the world (Conservation International 2021). Thus, these regions are among the most vulnerable ecosystems to the drivers of biodiversity loss. Anatolia (the Asian part of Turkey) is one of these regions because it is the region where three of these biodiversity hotspots meet, and interact: the (1) Caucasus, (2) Irano-Anatolian, and (3) Mediterranean Basin biodiversity hotspots. This means that Anatolia has a high biodiversity and a high percentage of life found nowhere else in the world, but has lost most of its original natural vegetation. In other words, Anatolia is one of the world’s biologically richest and most threatened terrestrial regions (Conservation International 2021, see also Gür, 2016, 2017a,b, Özüdoğru et al. 2020).

Anatolian ground squirrels, *Spermophilus xanthoprymnus* (Bennett 1835), are group-living, diurnal, hibernating, and pre-dominantly herbivorous, burrowing ground-dwelling squirrels (Kart Gür and Gür 2010). Anatolian ground squirrels hibernate individually in underground burrows, about from late summer to early spring (Gür and Kart Gür 2005, Kart Gür et al. 2009, Kart Gür and Gür 2010, 2015, 2017). Anatolian ground squirrels range about from 800 to 2,900 m and inhabit steppe or anthropogenic steppe areas (for detailed information on the Anatolian steppes, see FAO-TOB 2020) throughout the central and eastern Anatolia, with minor range extensions into the western Armenia and northwestern Iran, but do not inhabit large steppe areas generally < 1000 m in the southeastern Anatolia (Kryštufek and Vohralík 2005, 2012, Kart Gür and Gür 2010, Gür 2013). So, Anatolian ground squirrels are distributed mainly in the Anatolian section of the Irano-Anatolian biodiversity hotspot, excluding its middleeastern Anatolian part (Kart Gür and Gür 2017, Figure 1). The Irano-Anatolian biodiversity hotspot contains 2,500 endemic plant species and retains 3.6% of hotspot area as natural intact vegetation (Sloan et al. 2014, Habel et al. 2019). By the year 2050, it is projected to experience a significant change in climate for 15-24% of hotspot area and to lose all accessible natural intact vegetation to agriculture and therefore all endemic plant species (Habel et al. 2019). Already, Anatolian steppes, all located in the Irano-Turanian phytogeographical region, are the most threatened ecosystems among all natural ecosystems due to human activities (FAO-TOB 2020).

**Figure 1.**
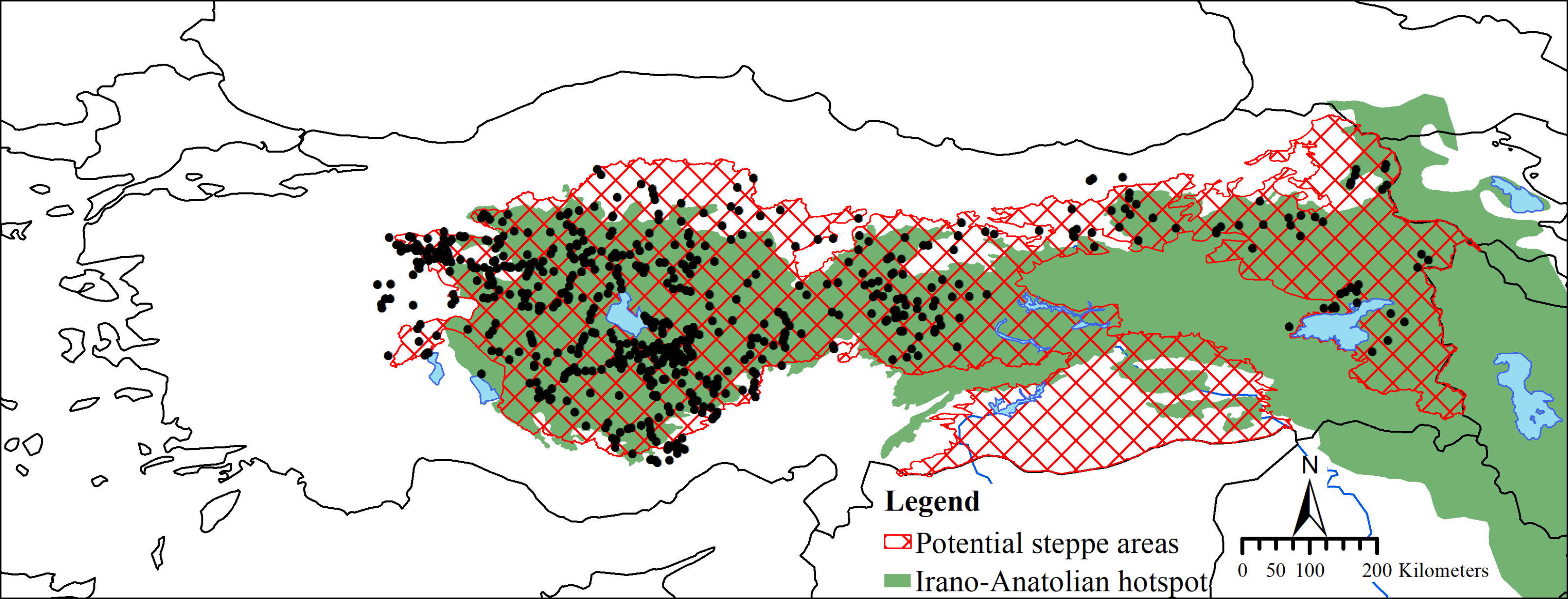
The geographic distribution (all presence records from the Monitoring Project for the Effects of Environmental Changes on Ground Squirrels, black circles) of Anatolian ground squirrels (*Spermophilus xanthoprymnus*), with potential steppe areas in Anatolia (the Nature Conservation Center, DKM, and Ambarlı et al. 2016) and the Irano-Anatolian hotspot in the study area (Conservation International 2021). The visible area in maps is 25° to 46°E and 35° to 43°N.

Anatolian ground squirrels are considered as ‘Near Threatened’ “due to population declines estimated at 20-25% over the last 10 years as a result of habitat conversion for agriculture, especially in central Anatolia” and also almost qualified as “Vulnerable” under criterion A2c because agriculture, a key part of Turkey’s economy, continues to be developed (Yiğit and Ferguson 2020). Contrary to that of most temperate (mid-latitude) species, Anatolian ground squirrels expanded rather than contracted their range during the glacial periods and contracted rather than expanded their range during the interglacial periods. According to this impact of the Late Quaternary glacial-interglacial cycles, ongoing climate change should also have been contributing to this population decline (Gür 2013). Thus, Anatolian ground squirrels are of most immediate concern because they currently contract their range and loss their habitats and therefore face increased threat with further increases in global temperature and land-use.

In this study, the future impact of climate and land-use changes on Anatolian ground squirrels was assessed. Accordingly, following the InSiGHTS modelling framework (Baisero et al. 2020), a hierarchical approach with two steps was used. First, ecological niche modelling, widely used to predict the impact of climate change on the distribution patterns of species (Franklin 2010, Peterson et al. 2011), with the presence records (from the project directed by the author) and bioclimatic data (from the WorldClim database for the present and future, Fick and Hijmans 2017), was used to assess the impact of climate change in areas accessible to Anatolian ground squirrels through dispersal (i.e. the impact of climate change). Gür (2013) suggests that climate is one of the main factors that limit the geographical distribution of Anatolian ground squirrels, and therefore they represent an ideal study system for niche modelling. Second, based on the habitat preferences of ground squirrels (Kryštufek and Vohralík 2005, 2012, Kart Gür and Gür 2010, Thorington et al. 2012), land-use data (from the Land-Use Harmonization dataset for the present and future, Hurtt et al. 2020) were used to assess the impact of land-use change in suitable bioclimatic areas for Anatolian ground squirrels under present and future conditions (i.e. the combined impact of both changes). Also, priority areas for the conservation of Anatolian ground squirrels were identified based on *in-situ* climate change refugia.

## 2. Methods

### 2.1. Ecological niche modelling

For Anatolian ground squirrels (*Spermophilus xanthoprymnus*), presence data (649 records, Figure 1) were obtained from the Monitoring Project for the Effects of Environmental Changes on Ground Squirrels that also gathers presence records for ground squirrels living in Anatolia, mostly by means of field studies (63 of these data from Gündüz et al. 2007, 127 from scientists and citizen scientists and validated by the author, and 459 from field studies made by the author as a part of the monitoring project). All these records have coordinates and therefore were used “as is”. The date range of these records spanned 1990s-2019 (mostly on 2010-2019). To reduce the effects of spatial sampling biases (Boria et al. 2014), the presence records were spatially filtered as follows, using the software SDMtoolbox v2.4 (Brown et al. 2017). First, environmental heterogeneity was calculated using a moving window approach (5×5 pixels of 2.5 arc-minutes) and the first three principal components of bioclimatic variables (excluding BIO8, 9, 18, and 19 because of known spatial artefacts, i.e. artificial discontinuities in climate gradients; for bioclimatic variables, see below), and reclassified into three broad classes (representing low, medium, and high heterogeneity) using the equal area classification type. Then, the presence records were spatially filtered by reducing multiple records to a single record within 5, 10, and 15 km distances in areas of high, medium, and low environmental heterogeneity, respectively, resulting in 285 presence records (Figure 2).

**Figure 2.**
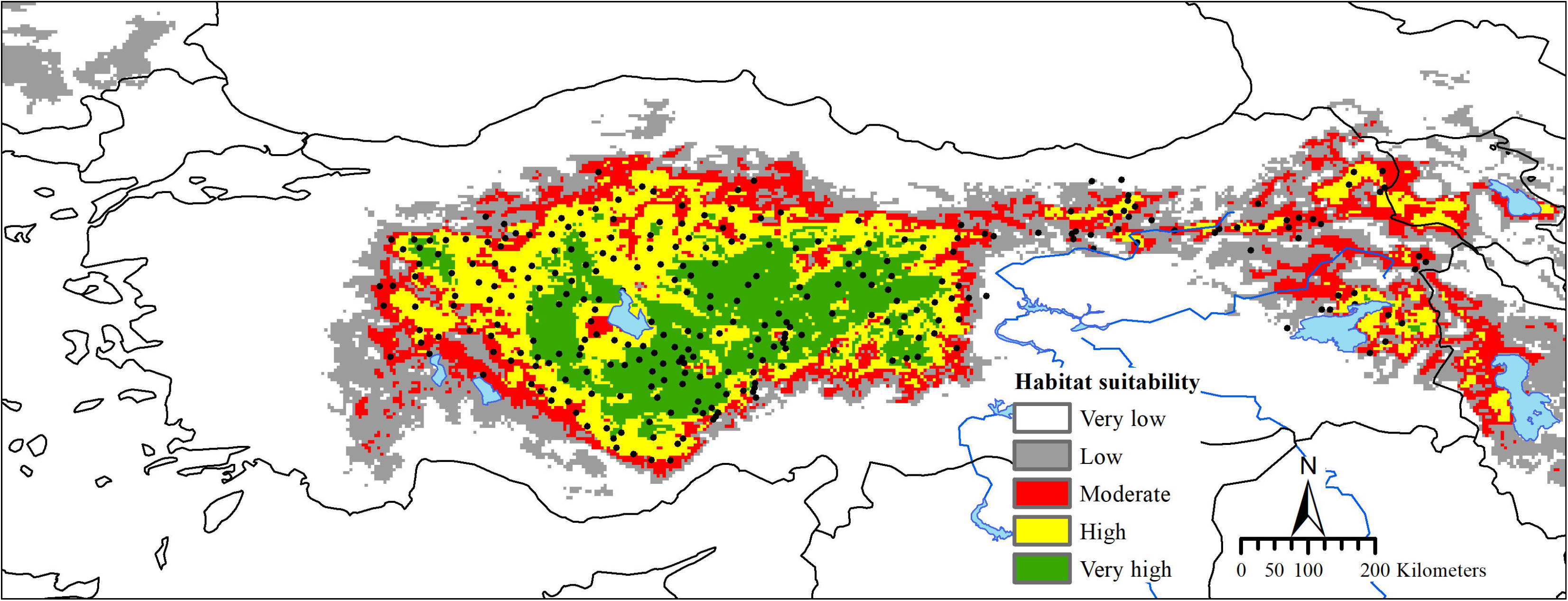
Habitat suitability under present (1970-2000) conditions for Anatolian ground squirrels (*Spermophilus xanthoprymnus*). Black circles indicate 285 presence records.

Bioclimatic data were downloaded from the WorldClim database version 2.1 (Fick and Hijmans 2017) at a spatial resolution of 2.5 arc-minutes (~ 4.6 km at the equator) for the present (1970-2000) and future (2030, average from 2021 to 2040, and 2050, average from 2041 to 2060). Future bioclimatic data from the Coupled Model Intercomparison Project Phase 6 (CMIP6) are, in this study, based on four global climate models (BCC-CSM2-MR, CNRM-CM6-1, CanESM5, and MIROC6) and four coupled SSP(shared socio-economic pathways)/RCP(representative concentration pathways) scenarios (from the low to the medium to the high forcing levels: SSP1/RCP2.6 (low), SSP2/RCP4.5 (medium), SSP3/RCP7.0 (high), and SSP5/RCP8.5 (high); for detailed information, see the WorldClim database and Riahi et al. 2017). The use of different global climate models and scenarios allowed assessing uncertainty in ecological niche modelling due to a broad range of possible climate change futures. These data include 19 scenopoetic bioclimatic variables (not affected by the presence of the focal species, Peterson et al. 2011) derived from monthly temperature and precipitation values (for detailed descriptions of bioclimatic variables, see the WorldClim database and Supplemental Table 1; see also Booth et al. 2014). In addition to bioclimatic data, topographic data were downloaded from the EarthEnv project (Amatulli et al. 2018) at a spatial resolution of 5 km. These data include scenopoetic topographic continuous and categorical variables derived from digital elevation models (for detailed descriptions of topographic variables, see the EarthEnv project and Amatulli et al. 2018). Based on what is known about the ecology of ground squirrels (Gür 2010, 2013, Kart Gür and Gür 2010, Gür and Kart Gür 2012, Gür et al. 2018) and preliminary analyses over years, three sets of non-collinear (r < □0.80□) variables were created to account for multicollinearity (for a similar approach, see Gür et al. 2018; Supplemental Table 1, 2 and Supplemental Figure 1). All these variables were masked to include Anatolia and surrounding areas (i.e. the study area, 25° to 46° E and 35° to 43° N, Gür 2013, Figure 1).

To model the Grinnellian niche (focusing on a species’ scenopoetic requirements, but not on the effects of the species on a given habitat, as considered by Charles Elton, Peterson et al. 2011, Anderson 2013) and to infer habitat suitability (or suitable bioclimatic areas) throughout the study area in the present and future for Anatolian ground squirrels, the software Maxent v3.4.1 (Phillips et al. 2017) was used because it requires presence-only data (not including any information on absence or non-detection, Peterson et al. 2011) and, in the circumstance of that data are presence-only format (as in this study), is a better choice over complicated, computationally intensive ‘black-box’ ensemble models (Kaky et al. 2020; for a supportive finding by using an ensemble approach, see Supplemental Box 1). The optimal set of variables and model settings (Elith et al. 2011, Merow et al. 2013) were selected in the software WALLACE v1.0.6.3, which “is an open-source GUI application that offers user-friendly access to R-scripted modern workflows” (Kass et al. 2018; for detailed methodological descriptions, see Muscarella et al. 2014), as follows. To select the background extent (or calibration area), a minimum convex polygon was created from the presence records to which a 2 degree buffer was applied (Supplemental Figure 2). This extent represents areas accessible to Anatolian ground squirrels through dispersal, termed M (Soberón and Peterson 2005). To ensure a full representation of environments available for Anatolian ground squirrels, following Guevara et al. (2018), all pixels within this extent was sampled as the background data (n = 55,079 pixels of 2.5 arc-minutes). To select the optimal set of variables and to regulate model complexity, 150 candidate models were tested by combining (a) three sets of non-collinear variables, (b) five combinations of feature classes (linear; linear and quadratic; hinge; linear, quadratic, and hinge; linear, quadratic, hinge, and product), and (c) ten values of regularization multiplier (1 to 5 in increments of 0.5). To calculate model evaluation statistics (for detailed methodological descriptions, see Muscarella et al. 2014, Kass et al. 2018 and Table 1), first, the presence and background data (i.e. the entire, unpartitioned dataset) were partitioned into separate training and testing bins for k□fold cross□validation, using a spatial approach (block, k = 4) that partitions data according to the latitudinal and longitudinal lines, resulting in four bins of (as far as possible) equal numbers (Supplemental Figure 2), which is especially relevant in cross-time or cross-space model transfer. Then, five models were tested for each candidate model: four out of these models were iteratively built using three out of four bins for model training and the withheld bin for model testing to calculate threshold-independent (AUC_TEST_ and AUC_DIFF_) and threshold-dependent (OR_MIN_, ‘Minimum Training Presence’ omission rate and OR_10_, 10% training omission rate) evaluation statistics (averaged over the iteration), and one using the entire, unpartitioned dataset (corresponding to all four bins) to calculate the Akaike information criterion corrected for small sample sizes (AICc). The optimal set of variables and model settings were selected based on the model with the highest performing in terms of overfitting and discrimination from among candidate models with delta AICc values of ≤2.

**Table 1.**
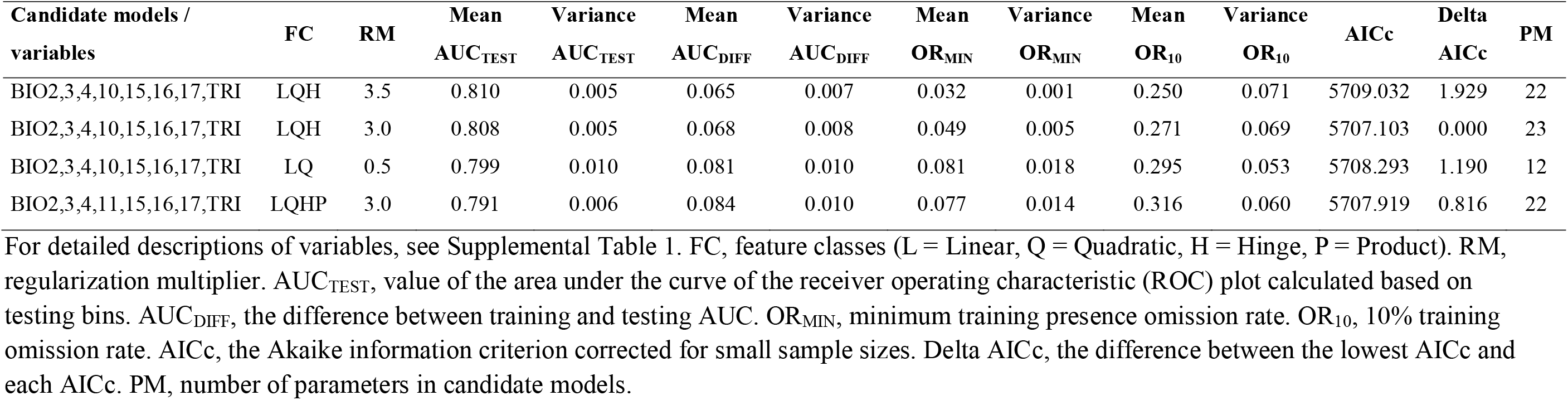
Model evaluation statistics of candidate models with delta AICc values of ≤2 ordered based on decreasing mean AUC_TEST_ value.

| Candidate models /<br>variables | FC | RM | Mean<br>AUC<sub>TEST</sub> | Variance<br>AUC<sub>TEST</sub> | Mean<br>AUC<sub>DIFF</sub> | Variance<br>AUC<sub>DIFF</sub> | Mean<br>OR<sub>MIN</sub> | Variance<br>OR<sub>MIN</sub> | Mean<br>OR<sub>10</sub> | Variance<br>OR<sub>10</sub> | AICc | Delta<br>AICc | PM |
| --- | --- | --- | --- | --- | --- | --- | --- | --- | --- | --- | --- | --- | --- |
| BIO2,3,4,10,15,16,17,TRI | LQH | 3.5 | 0.810 | 0.005 | 0.065 | 0.007 | 0.032 | 0.001 | 0.250 | 0.071 | 5709.032 | 1.929 | 22 |
| BIO2,3,4,10,15,16,17,TRI | LQH | 3.0 | 0.808 | 0.005 | 0.068 | 0.008 | 0.049 | 0.005 | 0.271 | 0.069 | 5707.103 | 0.000 | 23 |
| BIO2,3,4,10,15,16,17,TRI | LQ | 0.5 | 0.799 | 0.010 | 0.081 | 0.010 | 0.081 | 0.018 | 0.295 | 0.053 | 5708.293 | 1.190 | 12 |
| BIO2,3,4,11,15,16,17,TRI | LQHP | 3.0 | 0.791 | 0.006 | 0.084 | 0.010 | 0.077 | 0.014 | 0.316 | 0.060 | 5707.919 | 0.816 | 22 |

After the optimal set of variables and model settings were determined, a final model was developed using the entire, unpartitioned dataset (i.e. 285 presence records and 55 079 background points) and projected onto present and future conditions for the study area, with three options of extrapolation: unconstrained extrapolation, extrapolation with clamping, and no extrapolation (but only the results of unconstrained extrapolation are presented because the results of three options of extrapolation were similar, suggesting that extrapolation is not a critical issue). Multivariate environmental similarity surface (MESS) analysis (Elith et al. 2010) was performed to identify analog (similar) and non-analog (novel) conditions and extrapolation risks in model transfers. This analysis also provided information about which variable is driving novel conditions at any given pixel (i.e. the most dissimilar variable, Elith et al. 2010). Response curves (corresponding to a graph of the species’ response (here, habitat suitability) on the y-axis, versus the variable itself on the x-axis, Anderson 2013) were used to evaluate how each variable affects the prediction. Relative contribution of each variable to the final model was determined using a jackknife test. Cloglog output (ranging from 0 to 1, with 0 indicating low and 1 indicating high suitability) was selected to map habitat suitability (Phillips et al. 2017). Explain tool from Maxent (Elith et al. 2010) was used to investigate which variables are driving habitat suitability at any given pixel in the study area. Model performance and significance were evaluated by a partial ROC analysis (Peterson et al. 2008), as implemented in the software ntbox (NicheToolBox) v0.6.0.1 (Osorio-Olvera et al. 2020). This analysis was run with the following settings: proportion of omission = 0.05, percentage of random points = 50, and number of iterations for the bootstrap = 1,000. Also, response curves were examined if they are biologically logical.

To interpret easily, habitat suitability maps were categorized into five classes: very low suitability (< 0.2), low suitability (0.2-0.4), moderate suitability (0.4-0.6), high suitability (0.6-0.8), and very high suitability (> 0.8). Areas of moderate, high, very high suitability were also defined as suitable bioclimatic areas based on the ‘10 percentile training presence’ threshold (= 0.409). To reduce the number of maps and to facilitate the explanation of the results, only one map for each scenario (SSP1/RCP2.6, SSP2/RCP4.5, SSP3/RCP7.0, and SSP5/RCP8.5) for each time period (2030 and 2050) in the future as an average of four global climate models was presented. Areas of expansion, no change, and contraction in suitable bioclimatic areas under future conditions were calculated using the software SDMtoolbox v2.4 (Brown et al. 2017). Areas predicted suitable both in the present and future (across 2030 and 2050 for each scenario) were classified as *in-situ* climate change refugia (i.e. portions of a species’ present distribution projected to retain suitable conditions through time, Ashcroft 2010) in 2050 (Beaumont et al. 2019).

### 2.2. Land-use data

Land-use data were downloaded from the Land-Use Harmonization dataset version 2 (LUH2 v2f Release, Hurtt et al. 2020) at a spatial resolution of 0.25 degree (= 15.0 arc-minutes or ~ 28 km at the equator) for the present (2015) and future (2030 and 2050). This dataset provides the fraction of each of 12 land-use states for each cell, annually for the time period 850-2100. Land-use data from the Coupled Model Intercomparison Project Phase 6 (CMIP6) are, in this study, based on four coupled SSP/RCP scenarios (SSP1/RCP2.6 (sustainability-taking the green road, from IMAGE), SSP2/RCP4.5 (middle of the road, from MESSAGE-GLOBIOM), SSP3/RCP7.0 (regional rivalry-a rocky road, from AIM), and SSP5/RCP8.5 (fossil-fueled development-taking the highway, from REMIND-MAGPIE); for detailed information, see the LUH2 dataset and Riahi et al. 2017). The use of different scenarios allowed assessing uncertainty due to a broad range of possible land-use change futures. These data include 12 different land-use states: forested and non-forested primary and secondary lands, grazing lands (pasture and rangeland), urban land, and crops (C3 and C4 annual, C3 and C4 perennial, and C3 nitrogen-fixing).

Based on what is known about the habitat preferences of ground squirrels (Kryštufek and Vohralík 2005, 2012, Kart Gür and Gür 2010, Thorington et al. 2012) and the International Union for Conservation of Nature (IUCN) habitat classification scheme (Habitats: Grassland, Yiğit and Ferguson 2020) and also following Powers and Jetz (2019), non-forested primary and secondary lands were defined as suitable land-uses (i.e. habitats) for Anatolian ground squirrels, respectively. To infer habitat availability in the present and future, following the InSiGHTS modelling framework (Baisero et al. 2020), suitable habitats were calculated for each cell (i.e. the total fractions of non-forested primary and secondary lands x a layer cell area, km^2^) across suitable bioclimatic areas under present and future conditions, respectively. This approach allowed assessing the combined impact of climate and land-use changes. To interpret easily, habitat availability maps were categorized into five classes: very low availability (< 125 km^2^), low availability (125-250 km^2^), moderate availability (250-375 km^2^), high availability (375-500 km^2^), and very high availability (> 500 km^2^, note that, in the study area, the area of a grid cell is ~ 600 km^2^).

All GIS operations were conducted using the software ArcGIS v10.2.2.

## 3. Results

150 candidate models were tested based on different sets of non-collinear variables and model settings. The final model was developed using input variables of BIO2, 3, 4, 10, 15, 16, 17, and TRI; feature classes of linear, quadratic, and hinge; and a regularization multiplier of 3.5 (Table 1). This model used all of eight input variables (Supplemental Figure 3) and performed better than a random prediction (statistics for AUC ratio, mean ± SD = 1.73 ± 0.02, range = 1.67-1.77, P < 0.001).

Univariate response curves were bell-shaped for almost all of input variables and, if any, truncated when habitat suitability is decreasing and low (Supplemental Figure 4), suggesting that the background extent (or calibration area) contains the full range of scenopoetic conditions that Anatolian ground squirrels can inhabit (at least for input variables) or that the ‘niche space assumption’ (Anderson 2013) is met. Marginal response curves indicated that Anatolian ground squirrels primarily inhabit flat areas with colder, drier, and more seasonal bioclimatic conditions (Supplemental Figure 5, see also Supplemental Table 3). However, the variables that most contributed to the final model and therefore most influenced the geographic distribution of Anatolian ground squirrels were mean temperature of the warmest quarter (i.e. summer temperature), BIO10 (having the most useful information that is not present in the other variables) and precipitation of the wettest quarter, BIO16 (having the most useful information by itself) (Supplemental Figure 6). Accordingly, judging by corresponding response curves (Supplemental Figure 5), habitat suitability decreased with increasing summer temperature (especially > 20°C) and precipitation in the wettest quarter (generally occurring during spring with low precipitation in the central and northeastern Anatolia, and during autumn or winter with high precipitation in the other parts of Anatolia) (see Supplemental Figure 1).

Under present conditions, suitable bioclimatic areas (i.e. areas of moderate, high, very high suitability, 255,913 km^2^, 21.4% of the study area) were predicted mainly throughout the central and northeastern (Erzurum-Kars Plato and the north and east of Lake Van) Anatolia and adjacent Armenia and Iran (i.e. the Anatolian section of the Irano-Anatolian biodiversity hotspot, excluding its middleeastern Anatolian part). All these areas largely matched the geographical distribution of Anatolian ground squirrels (Figure 1 and 2), suggesting that they are at equilibrium or very near to equilibrium with scenopoetic conditions (i.e. the equilibrium distribution) or that the ‘noise assumptions’ (Anderson 2013) are met. Accordingly, large steppe areas in the southeastern Anatolia were correctly predicted to have very low suitabilities (Figure 2), driven by high summer temperatures (Supplemental Figure 7).

Coupled SSP/RCP scenarios (averaged over global climate models) predict summer temperature 2.4-2.7°C and 2.9-4.5°C hotter and precipitation in the wettest quarter almost the same both in the background extent and suitable bioclimatic areas under present conditions in the future (2030 and 2050, respectively). Summer temperature increases are large for each scenario for each time period in the future, but the most drastic towards the high forcing level scenario (i.e. RCP8.5) and 2050 (Supplemental Table 3), especially for inner Anatolia. Under these future conditions, suitable bioclimatic areas were projected to decline by 32.8-41.6% (RCP7.0 on the low and RCP8.5 on the high end) and 46.5-76.4% (RCP2.6 on the low and RCP8.5 on the high end) in 2030 and 2050, respectively (Table 2, Figure 3, see also Supplemental Figure 8), mainly due to summer temperature increases (Supplemental Figure 9). The impact of climate change was highly drastic for each scenario for each time period in the future, but, as expected, the most drastic towards the highest forcing level scenario (i.e. RCP8.5) and 2050 (Table 2), with the greatest loss of suitable bioclimatic areas at lower elevations (Table 3) and in the western part of the central Anatolia and in the eastern Anatolia (Figure 3). Thus, *in-situ* climate change refugia (58,838-134,499 km^2^, 23.0-52.6% of suitable bioclimatic areas under present conditions, RCP8.5 on the low and RCP2.6 on the high end, Table 4) were projected mainly in the eastern and southeastern parts of the central Anatolia (Figure 4), suggesting these regions as priority areas for the conservation of Anatolian ground squirrels. These results are broadly agreed upon by all global climate models (Supplemental Figure 10). Moreover, novel conditions were located mainly in the southeastern part of the study area (Supplemental Figure 11), especially for summer temperature, further supporting that extrapolation is not a critical issue (see Methods).

**Figure 3.**
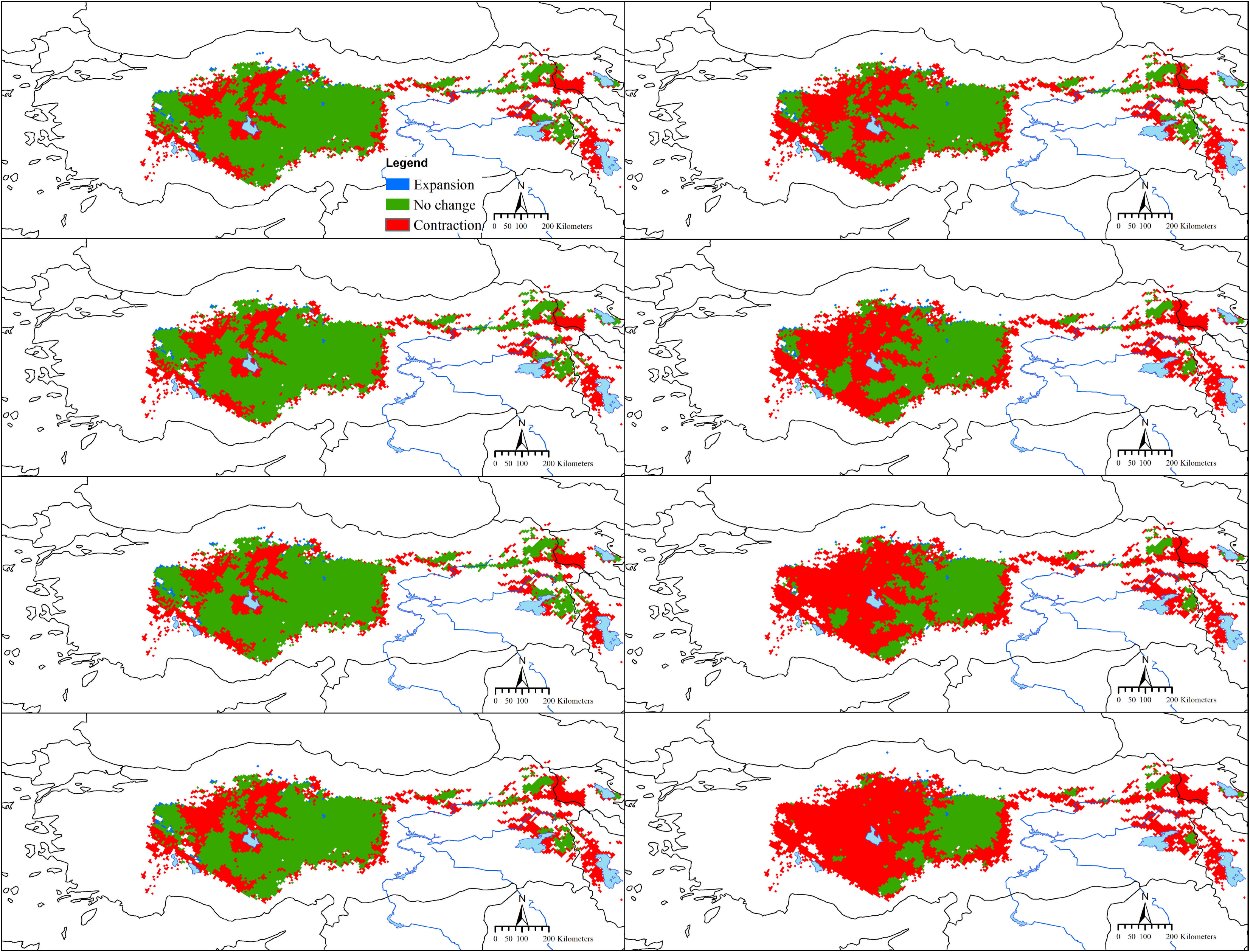
Areas of expansion, no change, and contraction in suitable bioclimatic areas under future conditions (i.e. for each scenario, SSP1/RCP2.6, SSP2/RCP4.5, SSP3/RCP7.0, and SSP5/RCP8.5, for each time period, 2030 and 2050) for Anatolian ground squirrels (*Spermophilus xanthoprymnus*). Left: 2030, right: 2050, from above to below: SSP1/RCP2.6, SSP2/RCP4.5, SSP3/RCP7.0, and SSP5/RCP8.5, respectively.

**Figure 4.**
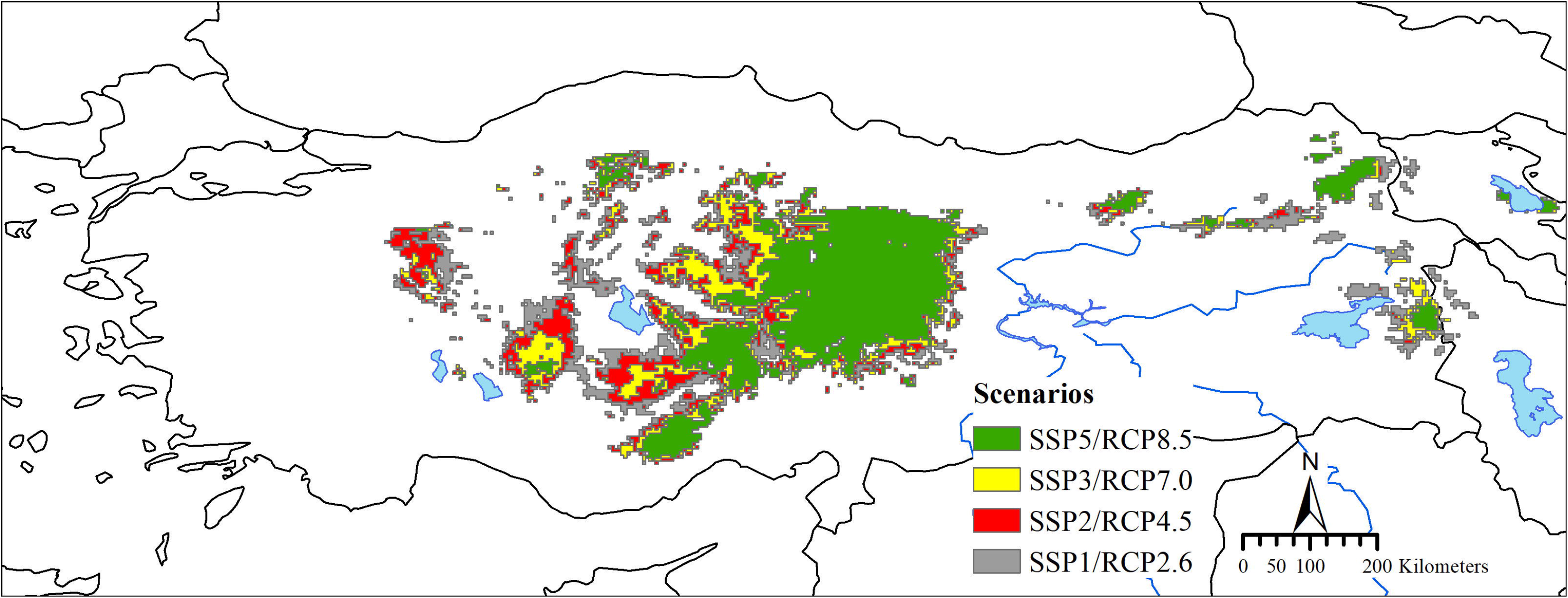
*In-situ* climate change refugia (i.e. areas predicted suitable both in the present and future, across 2030 and 2050 for each scenario, SSP1/RCP2.6, SSP2/RCP4.5, SSP3/RCP7.0, and SSP5/RCP8.5) for Anatolian ground squirrels (*Spermophilus xanthoprymnus*).

**Table 2.**
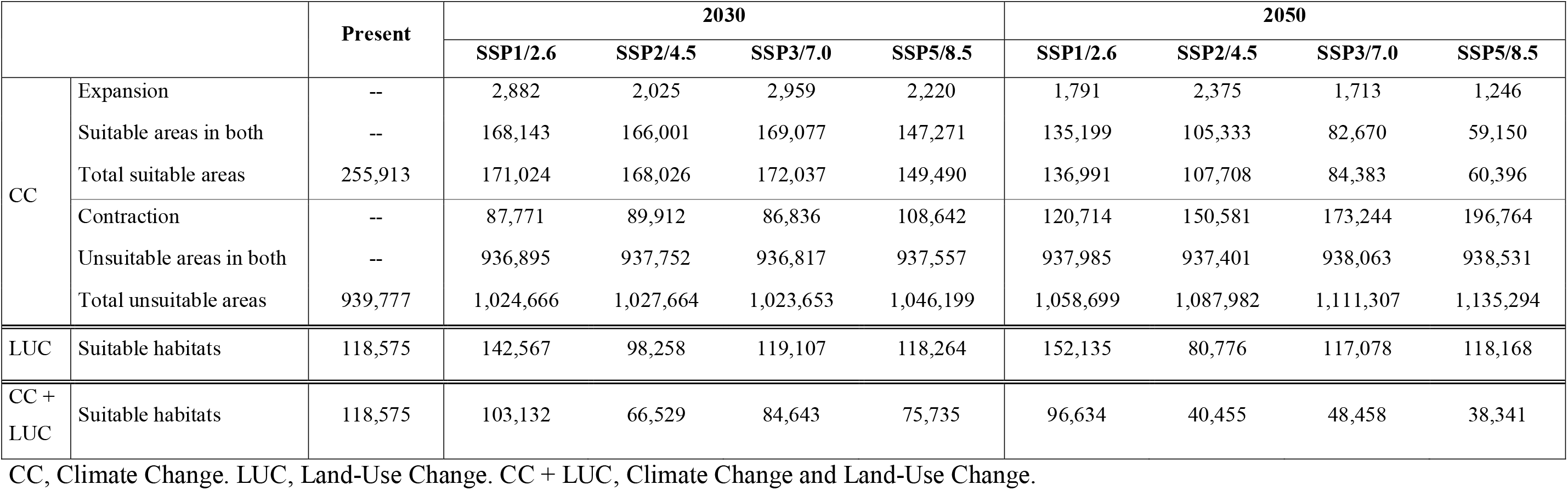
Habitat suitability (i.e. suitable bioclimatic areas, km^2^) and availability (i.e. suitable land-uses/habitats across suitable bioclimatic areas, km^2^) under present and future conditions (i.e. for each scenario, SSP1/RCP2.6, SSP2/RCP4.5, SSP3/RCP7.0, and SSP5/RCP8.5, for each time period, 2030 and 2050) for Anatolian ground squirrels (*Spermophilus xanthoprymnus*).

|  |  | Present | 2030 |  |  |  | 2050 |  |  |  |
| --- | --- | --- | --- | --- | --- | --- | --- | --- | --- | --- |
|  |  |  | SSP1/2.6 | SSP2/4.5 | SSP3/7.0 | SSP5/8.5 | SSP1/2.6 | SSP2/4.5 | SSP3/7.0 | SSP5/8.5 |
| CC | Expansion | -- | 2,882 | 2,025 | 2,959 | 2,220 | 1,791 | 2,375 | 1,713 | 1,246 |
|  | Suitable areas in both | -- | 168,143 | 166,001 | 169,077 | 147,271 | 135,199 | 105,333 | 82,670 | 59,150 |
|  | Total suitable areas | 255,913 | 171,024 | 168,026 | 172,037 | 149,490 | 136,991 | 107,708 | 84,383 | 60,396 |
|  | Contraction | -- | 87,771 | 89,912 | 86,836 | 108,642 | 120,714 | 150,581 | 173,244 | 196,764 |
|  | Unsuitable areas in both | -- | 936,895 | 937,752 | 936,817 | 937,557 | 937,985 | 937,401 | 938,063 | 938,531 |
|  | Total unsuitable areas | 939,777 | 1,024,666 | 1,027,664 | 1,023,653 | 1,046,199 | 1,058,699 | 1,087,982 | 1,111,307 | 1,135,294 |
| LUC | Suitable habitats | 118,575 | 142,567 | 98,258 | 119,107 | 118,264 | 152,135 | 80,776 | 117,078 | 118,168 |
| CC + LUC | Suitable habitats | 118,575 | 103,132 | 66,529 | 84,643 | 75,735 | 96,634 | 40,455 | 48,458 | 38,341 |

CC, Climate Change. LUC, Land-Use Change. CC + LUC, Climate Change and Land-Use Change.

**Table 3.** Median (interquartile range) elevation of suitable bioclimatic areas under present (1970-2000) and future conditions (i.e. for each scenario, SSP1/RCP2.6, SSP2/RCP4.5, SSP3/RCP7.0, and SSP5/RCP8.5, for each time period, 2030 and 2050).

| Present | 2030 |  |  |  | 2050 |  |  |  |
| --- | --- | --- | --- | --- | --- | --- | --- | --- |
|  | SSP1/2.6 | SSP2/4.5 | SSP3/7.0 | SSP5/8.5 | SSP1/2.6 | SSP2/4.5 | SSP3/7.0 | SSP5/8.5 |
| 1241 (1027-1616) | 1307 (1090-1639) | 1313 (1095-1644) | 1316 (1095-1650) | 1320 (1110-1631) | 1380 (1158-1691) | 1416 (1207-1693) | 1516 (1278-1742) | 1585 (1392-1787) |

**Table 4.** Habitat suitability (i.e. suitable bioclimatic areas, km^2^) and availability (i.e. suitable land-uses/habitats across suitable bioclimatic areas, km^2^) in *in-situ* climate change refugia for Anatolian ground squirrels (*Spermophilus xanthoprymnus*).

|  | Present | Climate Change Refugia |  |  |  |
| --- | --- | --- | --- | --- | --- |
|  |  | SSP1/2.6 | SSP2/4.5 | SSP3/7.0 | SSP5/8.5 |
| Habitat suitability | 255,913 | 134,499 | 104,787 | 82,436 | 58,838 |
| Habitat availability | 118,575 | 95,890 | 39,216 | 48,094 | 37,272 |

Suitable habitats were 46.3% (118,575 km^2^) of suitable bioclimatic areas under present conditions (Figure 5), suggesting that, as expected, land-use change already has a large impact on Anatolian ground squirrels. Habitat availability (i.e. suitable habitats across suitable bioclimatic areas) was projected to decline further by 13.0-43.9% (SSP1/RCP2.6 on the low and SSP2/RCP4.5 on the high end) and 18.5-67.7% (SSP1/RCP2.6 on the low and SSP5/RCP8.5 on the high end) in 2030 and 2050, respectively (Table 2, Figure 6). The combined impact of climate and land-use changes was highly drastic for each scenario for each time period in the future, but, as expected, the most drastic towards 2050, with marked differences between the sustainability/low forcing level scenario (i.e. SSP1/RCP2.6) and the other ones (Table 2, Figure 6). When the impact of climate and land-use changes was quantified separately, climate change appeared as the main driver of habitat loss because habitat availability were generally projected to remain almost same across suitable bioclimatic areas under present conditions in the future (2030 and 2050), except for the increase for SSP1/RCP2.6 and the decline for SSP2/RCP4.5 (Table 2). Suitable habitats within *in-situ* climate change refugia were 31.4-80.9% (37,272-95,890 km^2^, SSP5/RCP8.5 on the low and SSP1/RCP2.6 on the high end) of suitable habitats across suitable bioclimatic areas under present conditions (Table 4, Figure 7).

**Figure 5.**
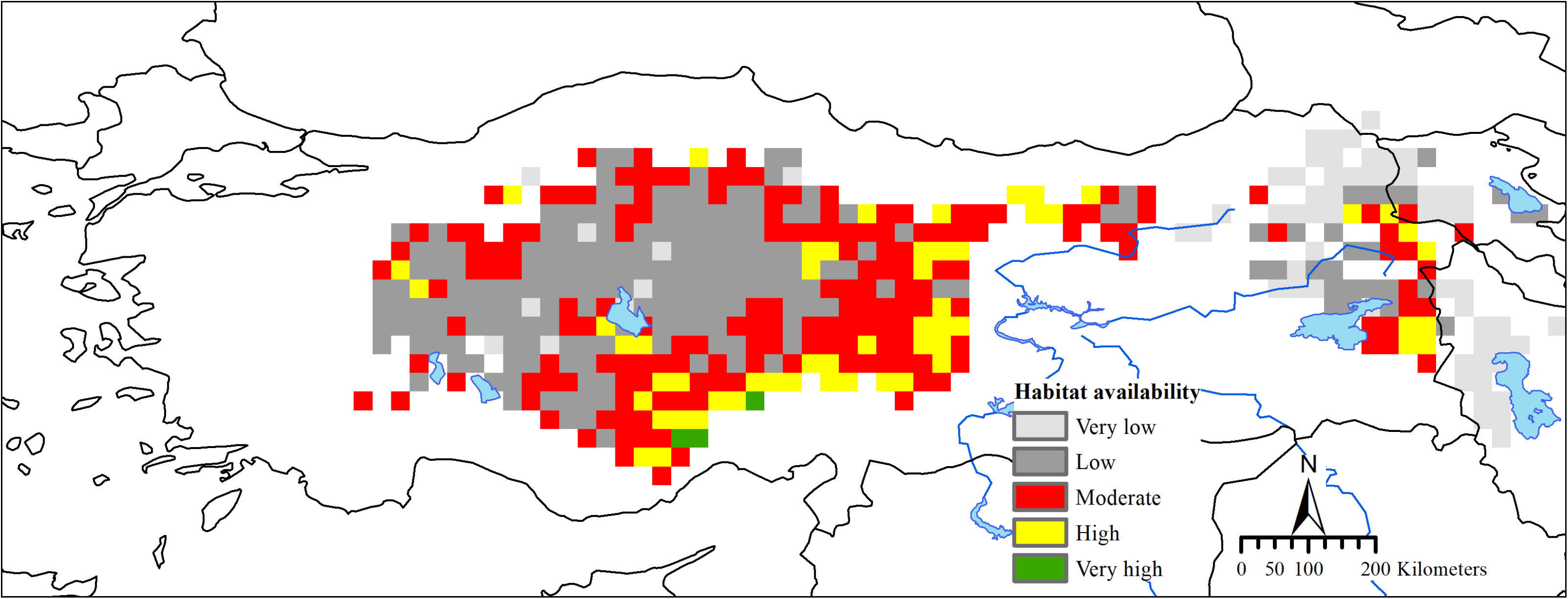
Habitat availability (i.e. suitable habitats across suitable bioclimatic areas) under present (1970-2000) conditions for Anatolian ground squirrels (*Spermophilus xanthoprymnus*).

**Figure 6.**
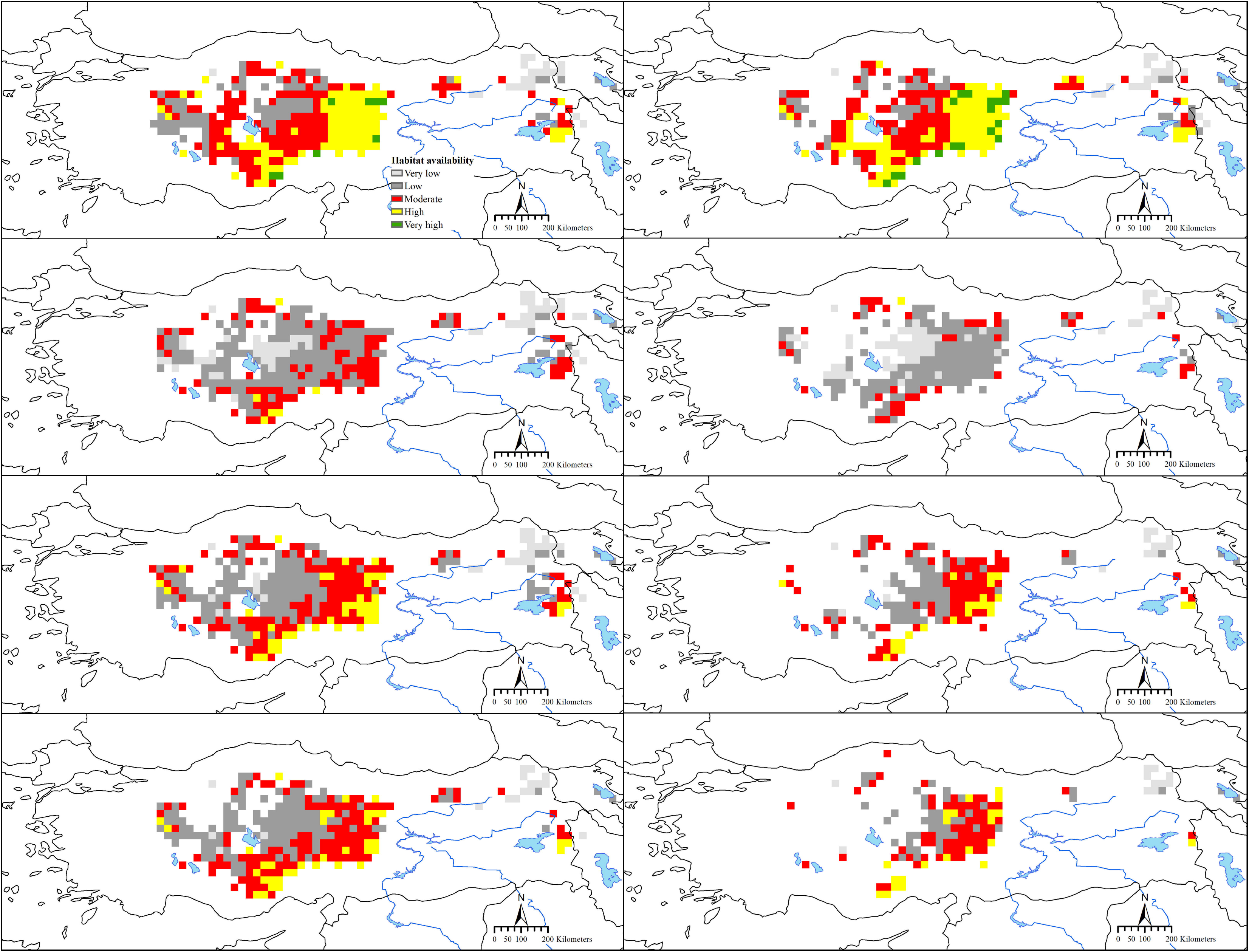
Habitat availability (i.e. suitable habitats across suitable bioclimatic areas) under future conditions (i.e. for each scenario, SSP1/RCP2.6, SSP2/RCP4.5, SSP3/RCP7.0, and SSP5/RCP8.5, for each time period, 2030 and 2050) for Anatolian ground squirrels (*Spermophilus xanthoprymnus*). Left: 2030, right: 2050, from above to below: SSP1/RCP2.6, SSP2/RCP4.5, SSP3/RCP7.0, and SSP5/RCP8.5, respectively.

**Figure 7.**
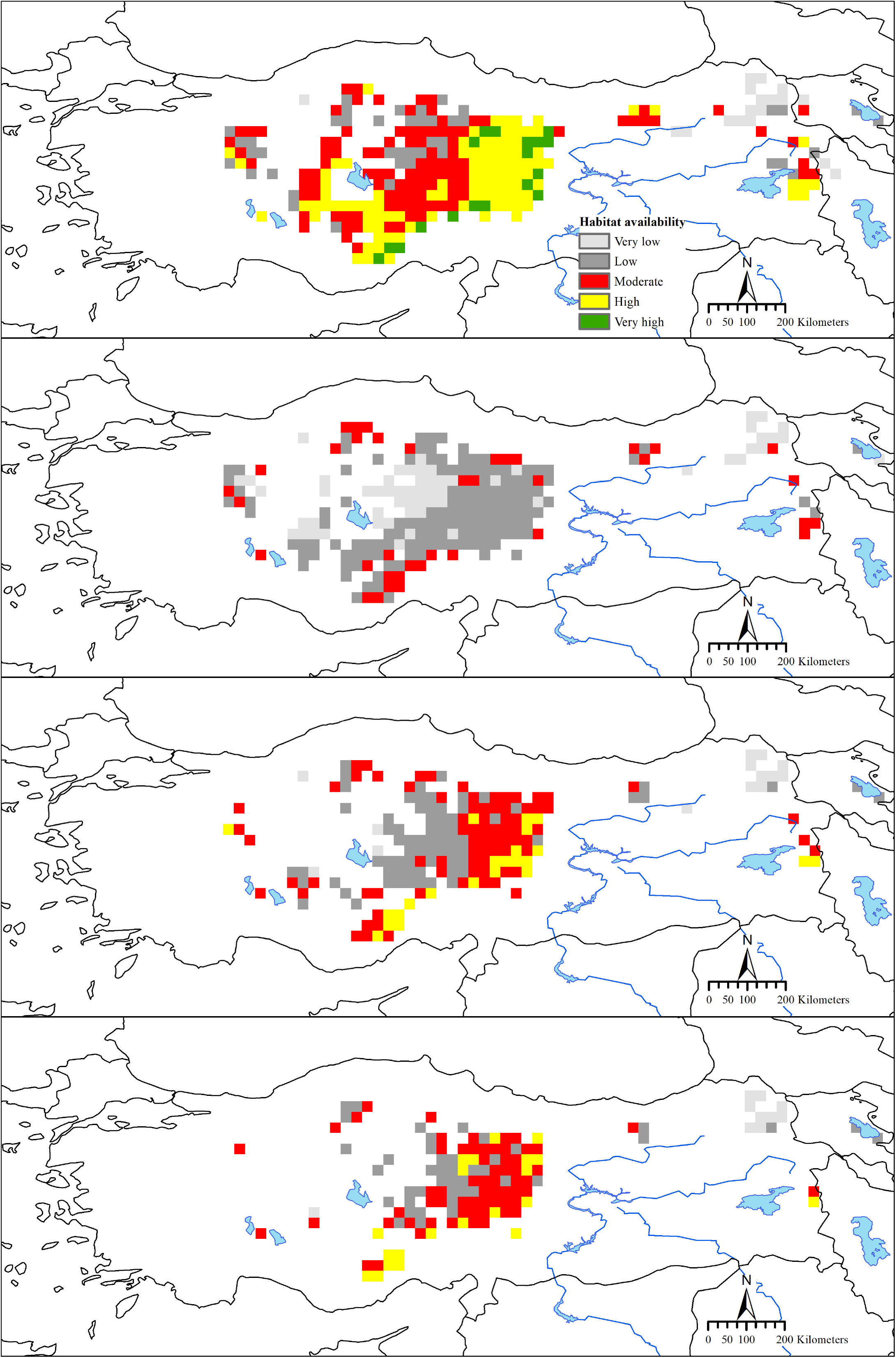
Habitat availability (i.e. suitable habitats across suitable bioclimatic areas) within *in-situ* climate change refugia (i.e. areas predicted suitable both in the present and future, across 2030 and 2050 for each scenario, SSP1/RCP2.6, SSP2/RCP4.5, SSP3/RCP7.0, and SSP5/RCP8.5) for Anatolian ground squirrels (*Spermophilus xanthoprymnus*). From above to below: SSP1/RCP2.6, SSP2/RCP4.5, SSP3/RCP7.0, and SSP5/RCP8.5, respectively.

## 4. Discussion

This study aimed to understand how climate and land-use changes will affect Anatolian ground squirrels in the future. Accordingly, following the InSiGHTS modelling framework (Baisero et al. 2020), a hierarchical approach with two steps was used: first, ecological niche modelling (Franklin 2010, Peterson et al. 2011) was used to infer suitable bioclimatic areas (or habitat suitability) throughout the study area in the present and future; and second, land-use data (Hurtt et al. 2020) were used to infer suitable habitats across suitable bioclimatic areas (i.e. habitat availability) under present and future conditions. This approach also allowed identifying priority areas for the conservation of Anatolian ground squirrels based on *in-situ* climate change refugia. This study represents a first attempt to combine niche modelling and land-use data for a species in Anatolia, which is one of the most vulnerable regions to the drivers of biodiversity loss (i.e. land-use change, direct exploitation of organisms, climate change, pollution, and invasion of alien species, IPBES 2019) because it is the region where three of biodiversity hotspots meet, and interact (Conservation International 2021).

Important advantage of this study is information about the distributional ecology of Anatolian ground squirrels that the Monitoring Project for the Effects of Environmental Changes on Ground Squirrels has provided. By July 2021, for Anatolian ground squirrels, over 1,200 presence records were collected, and the field routes of many thousands of kilometers were recorded by GPS. However, important limitation of this study is the temporal mismatch between the presence records of Anatolian ground squirrels and bioclimatic data. This is due to absence of the most recent bioclimatic data (also in terms of availability of the future CMIP data) or lack of the presence records from the recent past. Bioclimatic data were from the period 1970-2000, and the presence records were from the period 1990s-2019 (mostly after 2010). Such mismatch may be unavoidable for many of these kinds of studies, but the temporal range of presence and bioclimatic data should be matched whenever possible (Roubicek et al. 2010) Thus, future ecological niche modelling studies on Anatolian ground squirrels should also focus on this point.

Land-use change already has a large impact on Anatolian ground squirrels because suitable habitats were just 46% of suitable bioclimatic areas under present conditions. The International Union for Conservation of Nature (IUCN) Red List of Threatened Species estimates a 20-25% population decline over the last 10 years, due to large-scale agricultural activities that result in habitat loss and fragmentation, especially in the central Anatolia (Yiğit and Ferguson 2020). This is not really a surprise when considered that 77% of land (excluding Antarctica) has been modified by the direct effects of human activities (Pötrner et al. 2021), and at least 50% of the original vegetation in Anatolian steppes has been ploughed during the last century (Wesche et al. 2016). Moreover, Biodiversity Intactness Index indicates that land-use and related pressures have reduced local terrestrial biodiversity, especially in Anatolian steppes, over the recent years, 2000-2015 (Newbold et al. 2016, Supplemental Figure 12). Habitat availability was projected to decline further by 19-69% in the future (depending on the scenario), mainly due to the loss of suitable bioclimatic areas (47-77%, depending on the scenario) at lower elevations and in the western part of the central Anatolia and in the eastern Anatolia, suggesting that Anatolian ground squirrels will contract their range in the future, mainly due to climate change, as predicted by Gür (2013) (see also Introduction). All these are compatible with that land-use change is the greatest threat to biodiversity in the present (IPBES 2019), and climate change is likely to match or outpace land-use change in the future (Newbold 2018). By the year 2050, habitat availability is projected to decline by median 5-16% for 2,827 species of mammals, due to climate change rather than land-use change (Baisero et al. 2020). By the year 2100, at 1.5°C of global warming (above pre-industrial levels), 4% (2-7%) of 1,769 species of mammals are projected to lose over half of their climatically determined ranges, and at 4.5°C, this increases to 41% (29-57%) (Warren et al. 2018). Anatolian ground squirrels are one of these species, and therefore greater decline in habitat availability was projected for them than many species of mammals (19-69% vs. median 5-16%). However, it is important to note here that the approach in this study do not allow assessing the impact of habitat fragmentation (i.e. the breaking apart of habitats) on metapopulation dynamics. Thus, future ecological niche modelling and/or genetic studies on Anatolian ground squirrels should also focus on this point.

Hibernating species of ground squirrels (the genus *Spermophilus* sensu lato, Helgen et al. 2009) have a short active season, in which to reproduce and store fat reserves for survival during hibernation (Yensen and Sherman 2003, Thorington et al. 2012). Thus, much of the time spent above ground is allocated to foraging (Everts et al. 2004). However, thermal environment imposes substantial constraints on patterns of above ground activity and therefore can structure time available for foraging (Vispo and Bakken 1993, Sharpe and Van Horne 1999, Long et al. 2005) because ground squirrels are relatively small and inhabit generally open habitats (Yensen and Sherman 2003, Thorington et al. 2012), and therefore are subject to heat stress (Vispo and Bakken 1993, Sharpe and Van Horne 1999, Long et al. 2005). As other hibernating species of ground squirrels, Anatolian ground squirrels have a short active season, with individuals active above ground for mean 5.1-6.4 (the central Anatolia) and 3.9-4.7 (the northeastern Anatolia) months (Kart Gür et al. 2009, Kart Gür and Gür 2015, 2017). Individuals decrease aboveground activity at midday during hot days. In these days, individuals in traps sometimes cool themselves by using evaporative cooling of salivation and even die quite quickly, especially when traps are not checked regularly (Gür Kart and Gür 2010). These suggest that Anatolian ground squirrels are also very sensitive to overheating. Thus, warmer temperatures may reduce the time spent above ground for foraging (Vispo and Bakken 1993, Sharpe and Van Horne 1999, Long et al. 2005). In addition, global warming has increased and will increase agricultural and ecological droughts in the Mediterranean region (including Anatolian steppes), “which results from combined shortage of precipitation and excess evapotranspiration, and during the growing season impinges on crop production or ecosystem function in general” (IPCC 2021). All these, among other possible mechanisms, may influence life-history traits such as body mass, reproductive success, and survival of ground squirrels (Van Horne 2003). However, the approach in this study is correlation-based, which limits the inferences about possible mechanisms. Moreover, further limitations and caveats, among others (see Morin and Thuiller 2009, Ashcroft 2010, Morelli et al. 2016, Baumgartner et al. 2018, Beaumont et al. 2019), are that ecological niche modelling assumes that the present records represent viable populations and does not include any mechanistic relationships and take phenotypic plasticity and local adaptation into account. Thus, future studies (e.g. mechanistic niche modelling, ecophysiological ones etc.) on Anatolian ground squirrels should also focus on all these points.

Anatolian ground squirrels are highly sensitive to climate and land-use changes and therefore may be assessed to make inferences about these changes and monitored for early warnings of changes in steppe ecosystems (Siddig et al. 2016). Ground squirrels are keystone species, and therefore their loss may have serious consequences. For example, burrowing affects soil fertility, plant species composition, and primary productivity; encourages water infiltration; alters soil ion exchange capacity, water holding capacity, organic matter content, and inorganic nutrient levels; provides habitats for plants and animals, and increases grassland heterogeneity and landscape diversity (Yensen and Sherman 2003, Wesche et al. 2016). The present and future decline of Anatolian ground squirrels may threaten biodiversity in steppe areas because, for example, it may affect the specialized insects and produce a cascade effect on their predators. Anatolian ground squirrels are an important species in steppe areas for scarab dung beetle assemblages, with 12 ground squirrel-linked species (7 obligate + 5 associate), accounting for 15.6% of 77 species found in ground squirrel dens and livestock dung. Thus, population decline in Anatolian ground squirrels may cause a predictable decrease in the richness of dung beetle assemblages (Carpaneto et al. 2011). Anatolian ground squirrels are also an important prey species in steppe areas for a wide range of reptilian, avian, and mammalian predators (Kart Gür and Gür 2010). For example, breeding season diet of Saker Falcons (*Falco cherrug*) in Turkey typically comprises a greater proportion of Anatolian ground squirrels (61%, 35/57, of pellets and prey remains collected at three nest sites). Thus, population decline in Anatolian ground squirrels is identified as one of the main causes of decline in the number of Saker Falcon records in Turkey (Dixon et al. 2009). Thus, future studies on Anatolian ground squirrels should also focus on assessing their potential as an indicator species.

*In-situ* climate change refugia may facilitate the survival of existing populations and therefore the persistence of biodiversity especially during rapid climate change. Thus, one climate change adaptation strategy is to conserve these refugia to limit the impact of climate change in the 21st century (Morin and Thuiller 2009, Ashcroft 2010, Morelli et al. 2016, Baumgartner et al. 2018, Beaumont et al. 2019). Areas where these refugia are projected under most global climate models and coupled SSP/RCP scenarios may be sensible conservation targets because they are robust to uncertainty due to a broad range of possible climate change futures, leading to high stability in habitat suitability (Baumgartner et al. 2018). For Anatolian ground squirrels, these areas were projected mainly in the eastern and southeastern parts of the central Anatolia (especially in Kayseri, Nevşehir, Niğde, Sivas, and Yozgat provinces), in which habitat availability was also projected to be generally low and moderate, with some areas in the eastern Anatolia (Bayburt and Kars provinces). All these areas were generally located in the ‘Cool Temperate Dry’ zone of IPCC Climate Zones, i.e. in areas that are especially cooler than the surrounding region (Supplemental Figure 13), suggesting that many populations outside this zone (i.e. in the ‘Warm Temperate Dry’ zone) are very vulnerable to climate change, irrespective of global climate models and scenarios.

This will reduce overall genetic diversity because Anatolian ground squirrels are phylogeographically highly structured, with deep mitochondrial (mt) (sub)lineages restricted to small regions, or a large number of cytochrome *b* (cytb) mtDNA haplotypes (75%, 38 out of 51, based on data comprising 102 sequences from 70 localities) are restricted to a single locality (Gündüz et al. 2007, Harrison et al. 2003, Supplemental Figure 14). Only a very small part of Anatolian steppes (and therefore *in-situ* climate change refugia) lies within the protected areas (Supplemental Figure 15), majority of which are associated with wetlands, suggesting that the current protected areas are insufficient for the mitigation of climate and land-use change impacts on Anatolian steppes and their biodiversity (including Anatolian ground squirrels).

## Acknowledgments

I would like to acknowledge following scientists and citizen scientists for sharing their observations and participating in or being a guest in the field studies: E. Ada, S. Akkurt Gümüş, Ş. Aksan, E. Aktoklu, İ. B. Altıparmak, B. Aslan, S. Avcı, E. Aytekin, E. Berberoğlu, C. Bilgin, Ş. Bulut, L. Can, M. Çakır, M. Çelik, Ö. Çırpılı, H. Çohadar, Y. Demirbaş, A. Demirci, M. C. Demirok, A. Demirtaş, E. Dizdaroğlu, S. Ekşioğlu, C. Elverici, M. Elverici, G. Ergan, Ö. Erişöz Kasap, S. Ertuğrul, Y. Erzi, B. Fırat, M. Gaffaroğlu, D. Geçit, M. Gökmen, M. Göktuğ, A. Güngördü, Hilal Gür, S. Gür, İ. Güzel, S. Irmak, Teoman Kankılıç, Tolga Kankılıç, Ç. Karacaoğlu, E. Karaçetin, M. Karakaya, Adem Karataş, Ö. Keşaplı Can, S. T. Körüklü, Ü. Malkaçoğlu, D. Mengüllüoğlu, A. Öğretmen, E. Özay, O. Özbek, C. Özcan, M. Özcan, K. Özgişi, M. Z. Öztürk, B. Özüdoğru, M. Poyraz, D. Ragyov, Z. Sakarya, İ. K. Sağlam, A. Sefalı, H. Sevgili, L. Sınav, M. K. Şahin, N. Şalıkara, A. Şenel, F. M. Şimşek, B. Tatar, Ç. Tavşanoğlu, A. Tıraş, M. Toksoy, M. Topuz, TRAMEM, A. Turak, O. Ç. Türkay, Y. Uçarlı, N. Usta, E. D. Ülker, Z. Ünal, F. Ünsal, O. Ürker, B. Yalgın, S. Yanardağ, İ. Yıldırım, H. Yılmaz, and T. Yorulmaz, and especially M. Kart Gür. The Monitoring Project for the Effects of Environmental Changes on Ground Squirrels accelerated in 2018 with a series of field studies, while largely self-funded, did receive a small amount of support from the Kırşehir Ahi Evran University Scientific Research Projects Coordination Unit (Project Numbers: PYO-FEN.4001.12.012, completed in 2014 and PYO-FEN.4001.15.008, completed in 2018) and carried out by the permission of Republic of Turkey Ministry of Agriculture and Forestry, General Directorate of Nature Conservation and National Parks.

