## Supplemental File for "The future impact of climate and land-use changes on Anatolian ground squirrels under different scenarios"

**Supplemental Table 1.** Bioclimatic and topographic variables. Highlighted variables were used to create three sets of non-collinear variables.

| BIO1 | Annual Mean Temperature |
| --- | --- |
| BIO2 | Mean Diurnal Range |
| BIO3 | Isothermality |
| BIO4 | Temperature Seasonality |
| BIO5 | Max Temperature of Warmest Month |
| BIO6 | Min Temperature of Coldest Month |
| BIO7 | Temperature Annual Range |
| BIO8 | Mean Temperature of Wettest Quarter |
| BIO9 | Mean Temperature of Driest Quarter |
| BIO10 | Mean Temperature of Warmest Quarter |
| BIO11 | Mean Temperature of Coldest Quarter |
| BIO12 | Annual Precipitation |
| BIO13 | Precipitation of Wettest Month |
| BIO14 | Precipitation of Driest Month |
| BIO15 | Precipitation Seasonality |
| BIO16 | Precipitation of Wettest Quarter |
| BIO17 | Precipitation of Driest Quarter |
| BIO18 | Precipitation of Warmest Quarter |
| BIO19 | Precipitation of Coldest Quarter |
| TRI | Terrain Ruggedness Index |

**Supplemental Table 2.** Three sets of non-collinear (r < ǀ0.80ǀ) variables.

| Set I | BIO2 | BIO3 | BIO4 | BIO5 | BIO6 |  |  | BIO15 | BIO16 | BIO17 | TRI |
| --- | --- | --- | --- | --- | --- | --- | --- | --- | --- | --- | --- |
| Set II | BIO2 | BIO3 | BIO4 |  |  | BIO10 |  | BIO15 | BIO16 | BIO17 | TRI |
| Set III | BIO2 | BIO3 | BIO4 |  |  |  | BIO11 | BIO15 | BIO16 | BIO17 | TRI |

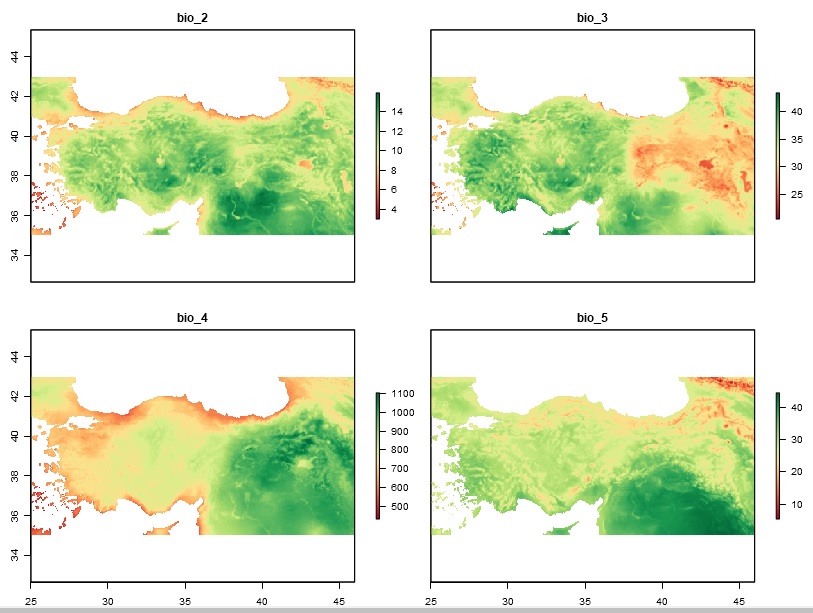

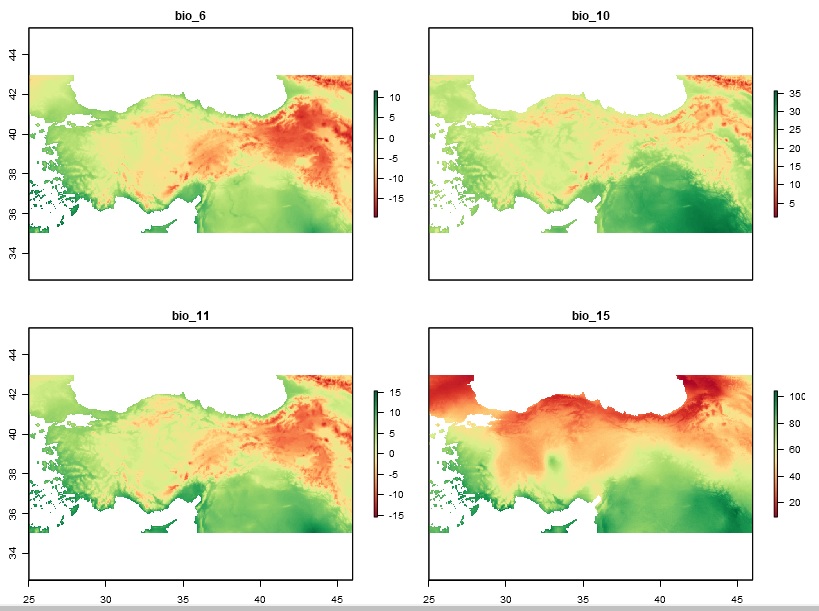

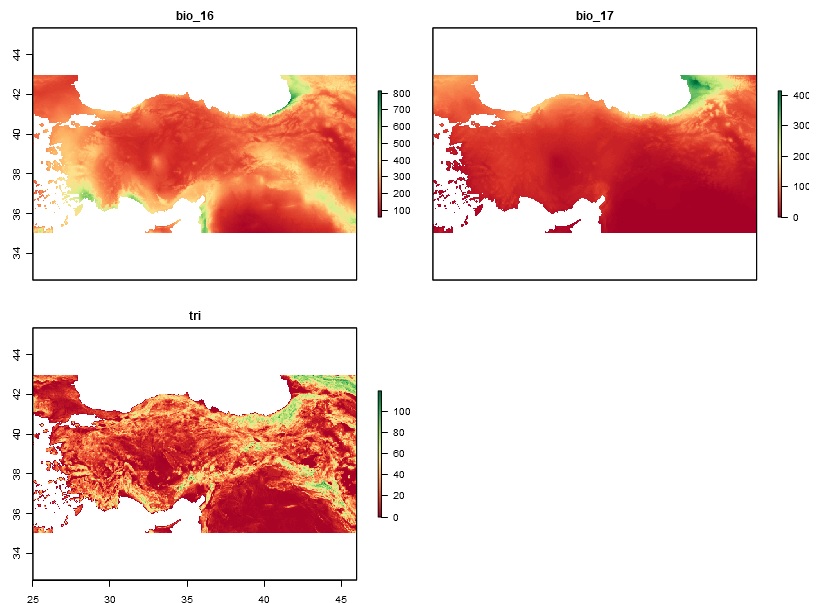

**Supplemental Figure 1.** Variables used to create three sets of non-collinear variables.

**Supplemental Box 1.** The similarity between the Maxent and ensemble predictions.

For the Maxent prediction, see the main text and below. The ensemble prediction for Anatolian ground squirrels (*Spermophilus xanthoprymnus*) was developed by the ‘SSDM’ R package (Schmitt et al. 2017) using the following data, algorithms, and default implementations.

**Occurrence and environmental data:** the same in the final model developed by the software Maxent.

**Algorithms:** artificial neural network (ANN), classification tree analysis (CTA), generalized additive model (GAM), generalized boosted regressions model (GBM), generalized linear model (GLM), multivariate adaptive regression splines (MARS), random forest (RF), and support vector machines (SVM)

**Repetitions:** 10

**Evaluation metric:** SES, which uses the sensitivity-specificity equality, as recommended by Liu et al. (2005)

**Number of and strategy used to select pseudo-absences:** automatic pseudo-absences with the recommendations from Barbet-Massin et al. (2012)

**Model evaluation method:** holdout with a fraction of 0.70

**Cross-validation repetitions:** 10

**Variable importance evaluation metric:** Pearson

**Ensemble weighted:** true

**Ensemble selection metric:** AUC, area under the receiver operating characteristic (ROC) curve

**AUC threshold:** 0.75

The similarity between the Maxent and ensemble predictions was calculated by ENMTools (Warren et al. 2010). ENMTools quantifies niche similarity using two similar metrics (*I* and Schoener’s *D*, Warren et al. 2008) that range from 0 (no similarity) to 1 (complete similarity).

All algorithms and the ensemble prediction performed better than a random prediction (AUC ≥ 0.773 and AUC = 0.831, respectively).

The variables that most contributed to the ensemble prediction were precipitation of the wettest quarter, BIO16 (variable relative contribution = 19.09%) and mean temperature of the warmest quarter (i.e. summer temperature), BIO10 (17.82%).

Maxent (see below) and the ensemble prediction performed similarly well (AUC = 0.863 and 0.831, respectively). The similarity between the MaxEnt and ensemble predictions was very high (*I* = 0.99 and Schoener’s *D* = 0.92).

Barbet‐Massin, M., Jiguet, F., Albert, C. H., Thuiller, W., 2012. Selecting pseudo‐absences for species distribution models: how, where and how many?. Methods in Ecology and Evolution 3, 327-338.

Liu, C., Berry, P. M., Dawson, T. P., Pearson, R. G., 2005. Selecting thresholds of occurrence in the prediction of species distributions. Ecography 28, 385-393.

Schmitt, S., Pouteau, R., Justeau, D., de Boissieu, F., Birnbaum, P., 2017. ssdm: An r package to predict distribution of species richness and composition based on stacked species distribution models. Methods in Ecology and Evolution 8, 1795-1803.

Warren, D. L., Glor, R. E., Turelli, M., 2008. Environmental niche equivalency versus conservatism: quantitative approaches to niche evolution. Evolution: International Journal of Organic Evolution 62, 2868-2883.

Warren, D. L., Glor, R. E., Turelli, M., 2010. ENMTools: a toolbox for comparative studies of environmental niche models. Ecography 33, 607-611.

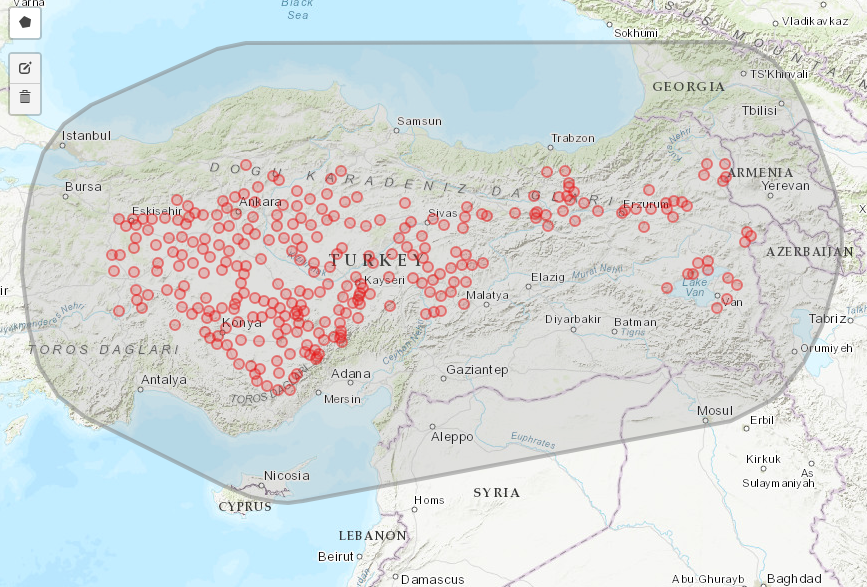

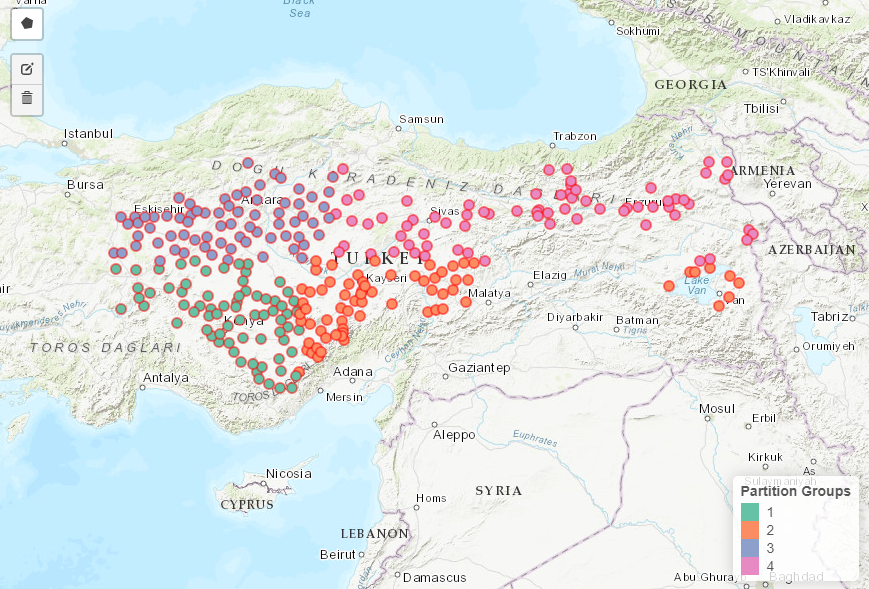

**Supplemental Figure 2.** (above) The minimum convex polygon created from the presence records (circles) to which a 2 degree buffer was applied and (below) the presence data (circles) partitioned into four bins of (as far as possible) equal numbers.

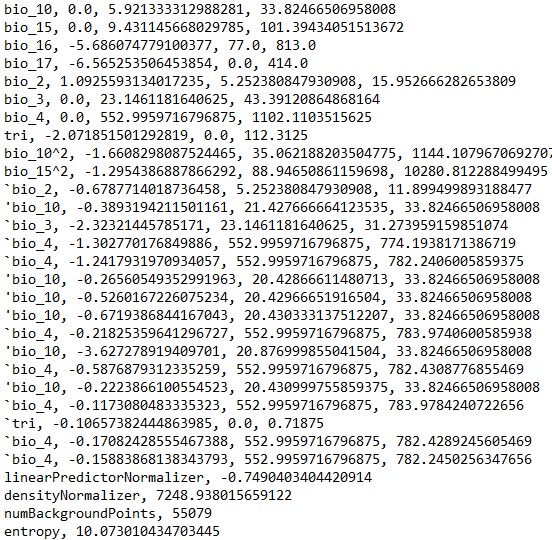

**Supplemental Figure 3.** The coefficients of the final model.

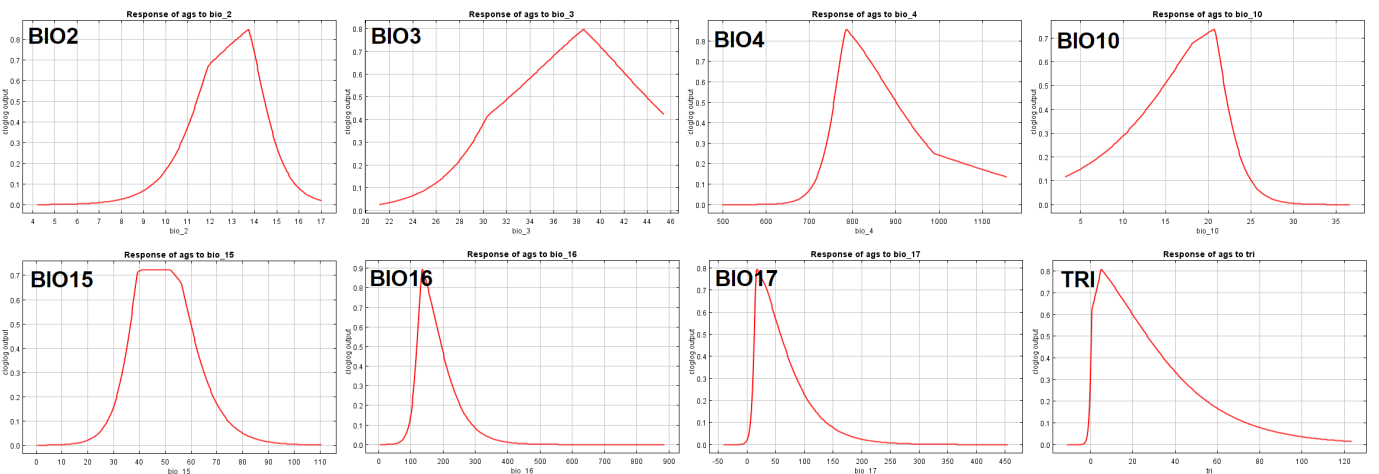

**Supplemental Figure 4.** The univariate response curves.

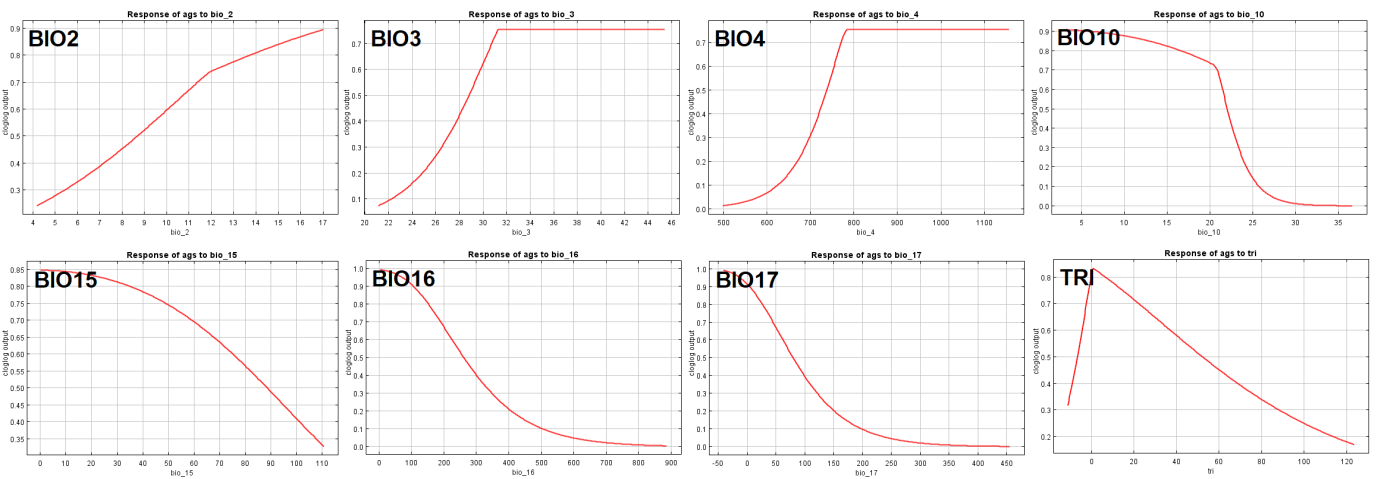

**Supplemental Figure 5.** The marginal response curves.

**Supplemental Table 3.** Medians and interguartile ranges of mean temperature of the warmest quarter (BIO10) and precipitation of the wettest quarter (BIO16) (averaged for each scenario, SSP1/RCP2.6, SSP2/RCP4.5, SSP3/RCP7.0, and SSP5/RCP8.5, over four global climate models, BCC-CSM2-MR, CNRM-CM6-1, CanESM5, and MIROC6) in the background extent (n = 10,000 pixels of 2.5 arc-minutes) and suitable areas (n = 2,770 pixels, note that these areas are under present, 1970-2000, conditions) both in the present and future (2030 and 2050).

|  |  | Present |
| --- | --- | --- |
| BIO10 | Suitable areas | 19.6 (18.0-20.6) |
|  | Background | 20.3 (17.9-23.7) |
| BIO16 | Suitable areas | 165 (145-187) |
|  | Background | 217 (172-283) |

|  |  | 2030 | | | |
| --- | --- | --- | --- | --- | --- |
|  |  | SSP1/RCP2.6 | SSP2/RCP4.5 | SSP3/RCP7.0 | SSP5/RCP8.5 |
| BIO10 | Suitable areas | 22.1 (20.5-23.1) | 22.0 (20.4-23.0) | 22.1 (20.5-23.1) | 22.3 (20.8-23.3) |
|  | Background | 22.7 (20.3-26.1) | 22.7 (20.3-26.0) | 22.8 (20.4-26.1) | 22.9 (20.6-26.4) |
| BIO16 | Suitable areas | 169 (149-190) | 170 (149-192) | 167 (147-188) | 169 (149-191) |
|  | Background | 220 (175-287) | 221 (176-288) | 216 (172-282) | 220 (175-285) |

|  |  | 2050 | | | |
| --- | --- | --- | --- | --- | --- |
|  |  | SSP1/RCP2.6 | SSP2/RCP4.5 | SSP3/RCP7.0 | SSP5/RCP8.5 |
| BIO10 | Suitable areas | 22.5 (21.0-23.6) | 23.1 (21.5-24.0) | 23.6 (22.0-24.6) | 24.1 (22.5-25.1) |
|  | Background | 23.2 (20.8-26.6) | 23.7 (21.3-27.1) | 24.2 (21.8-27.5) | 24.7 (22.2-28.0) |
| BIO16 | Suitable areas | 170 (150-192) | 167 (147-189) | 162 (142-183) | 165 (144-186) |
|  | Background | 222 (176-289) | 218 (173-283) | 213 (169-277) | 216 (171-281) |

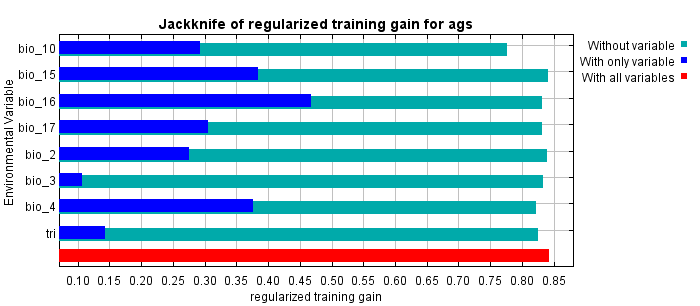

**Supplemental Figure 6.** The results of the jackknife test of variable importance.

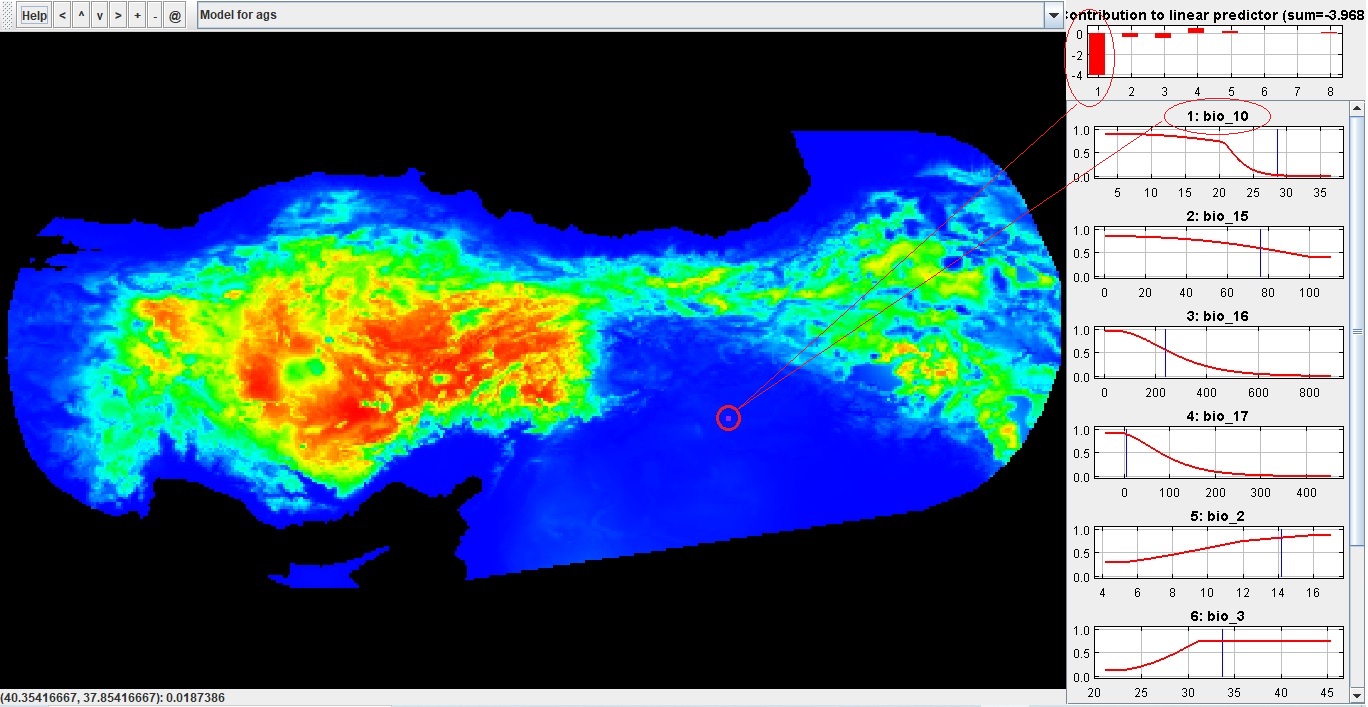

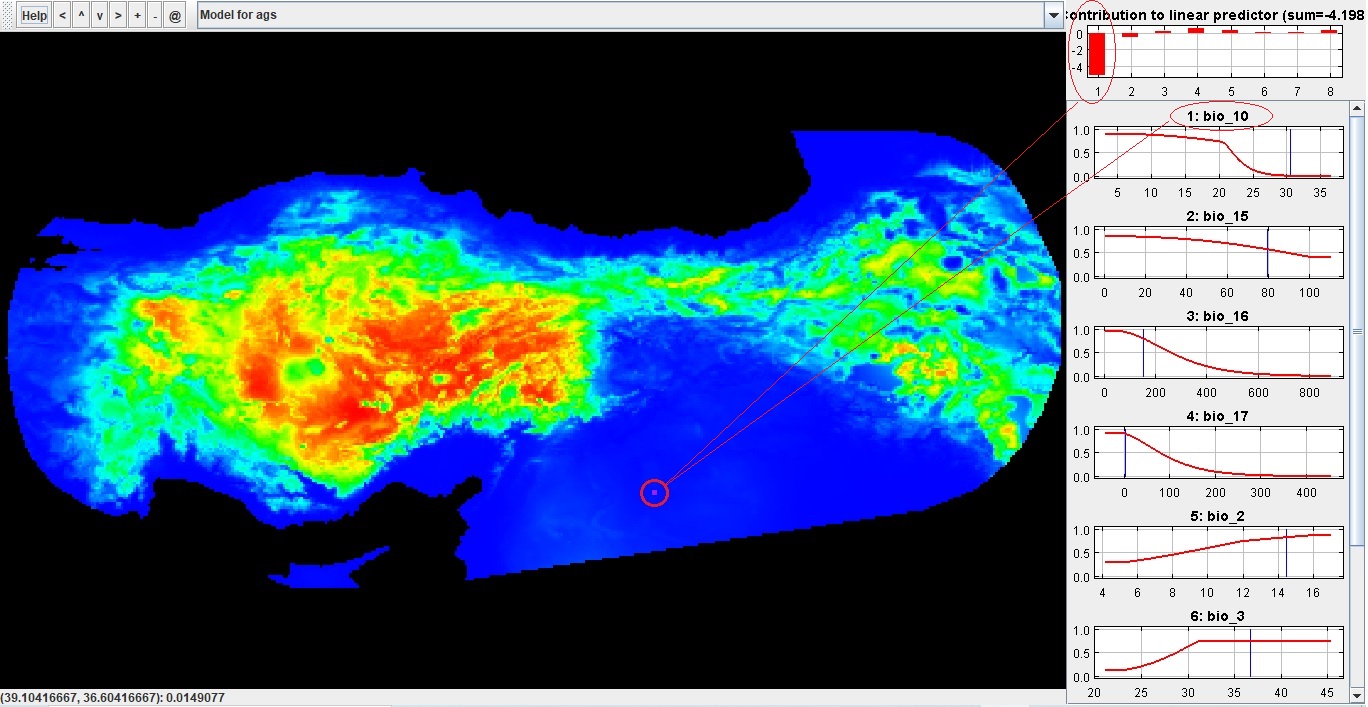

Supplemental Figure 7. Explain tool from Maxent. Warm colours represent high, and cold colours represent low habitat suitability under present (1970-2000) conditions. The effect of variables is explored at point locations in the southeastern Anatolia. Very low suitabilities in this region were driven by high summer temperatures (BIO10).

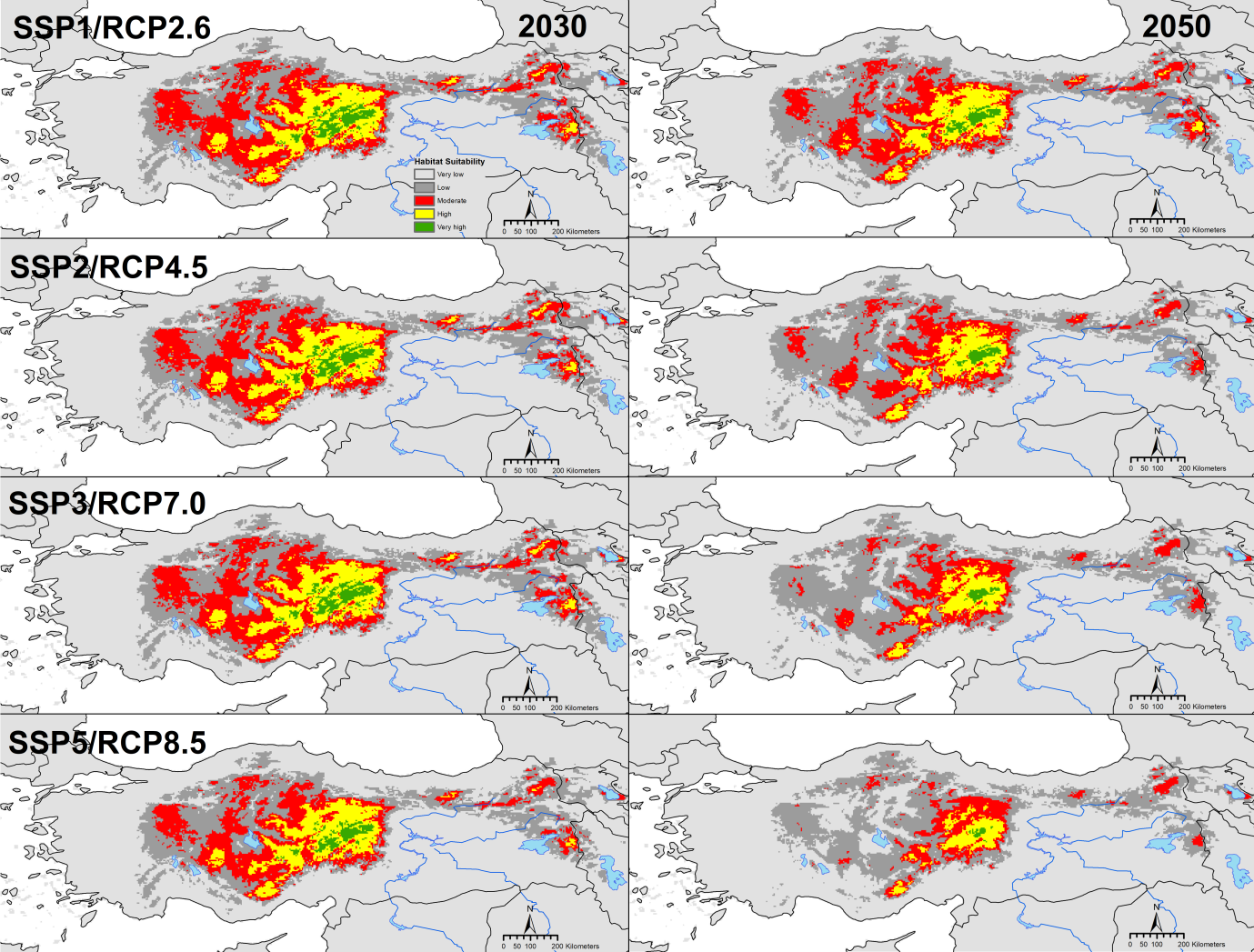

**Supplemental Figure 8.** Habitat suitability under future conditions (i.e. for each scenario, SSP1/RCP2.6, SSP2/RCP4.5, SSP3/RCP7.0, and SSP5/RCP8.5, for each time period, 2030 and 2050).

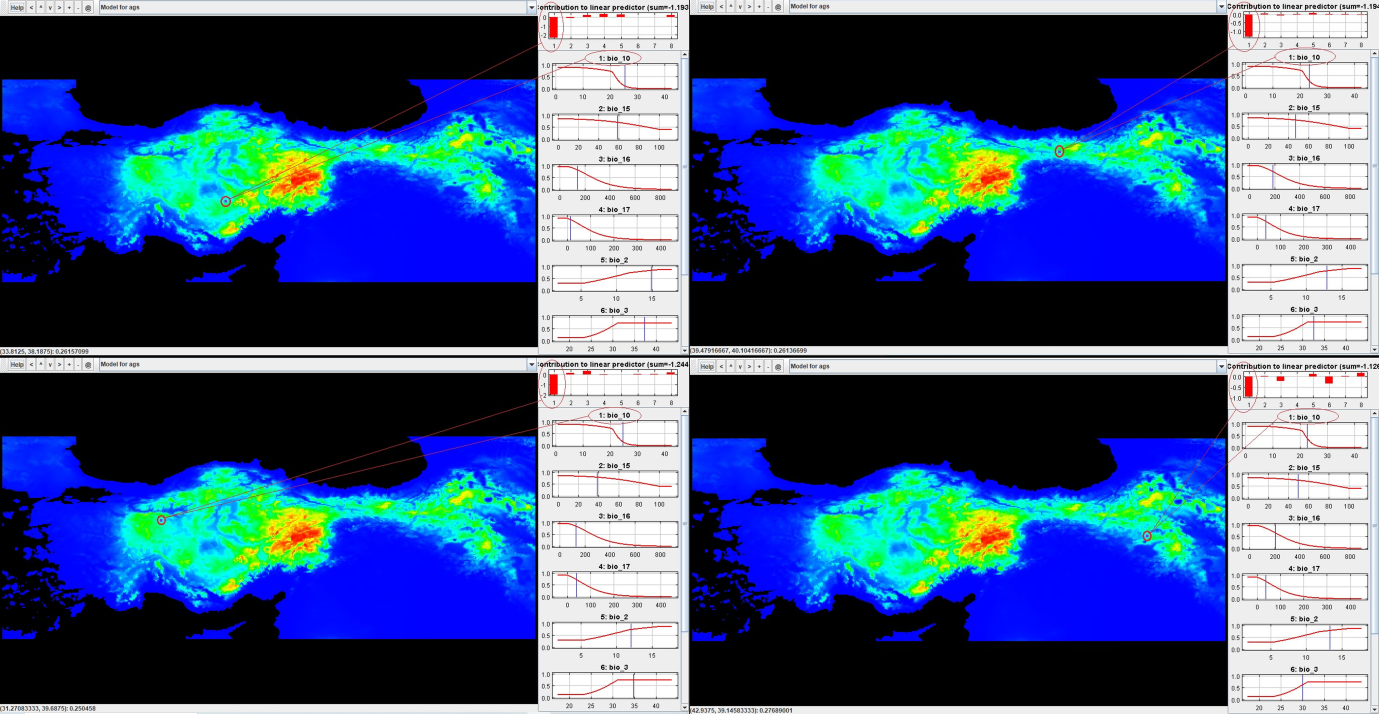

Supplemental Figure 9. Explain tool from Maxent. Warm colours represent high, and cold colours represent low habitat suitability under future conditions (e.g. for SSP5/RCP8.5 for 2050 for MIROC6). The effect of variables is explored at point locations in suitable bioclimatic areas under present conditions. Low suitabilities in these areas under future conditions were mainly driven by high summer temperatures (BIO10).

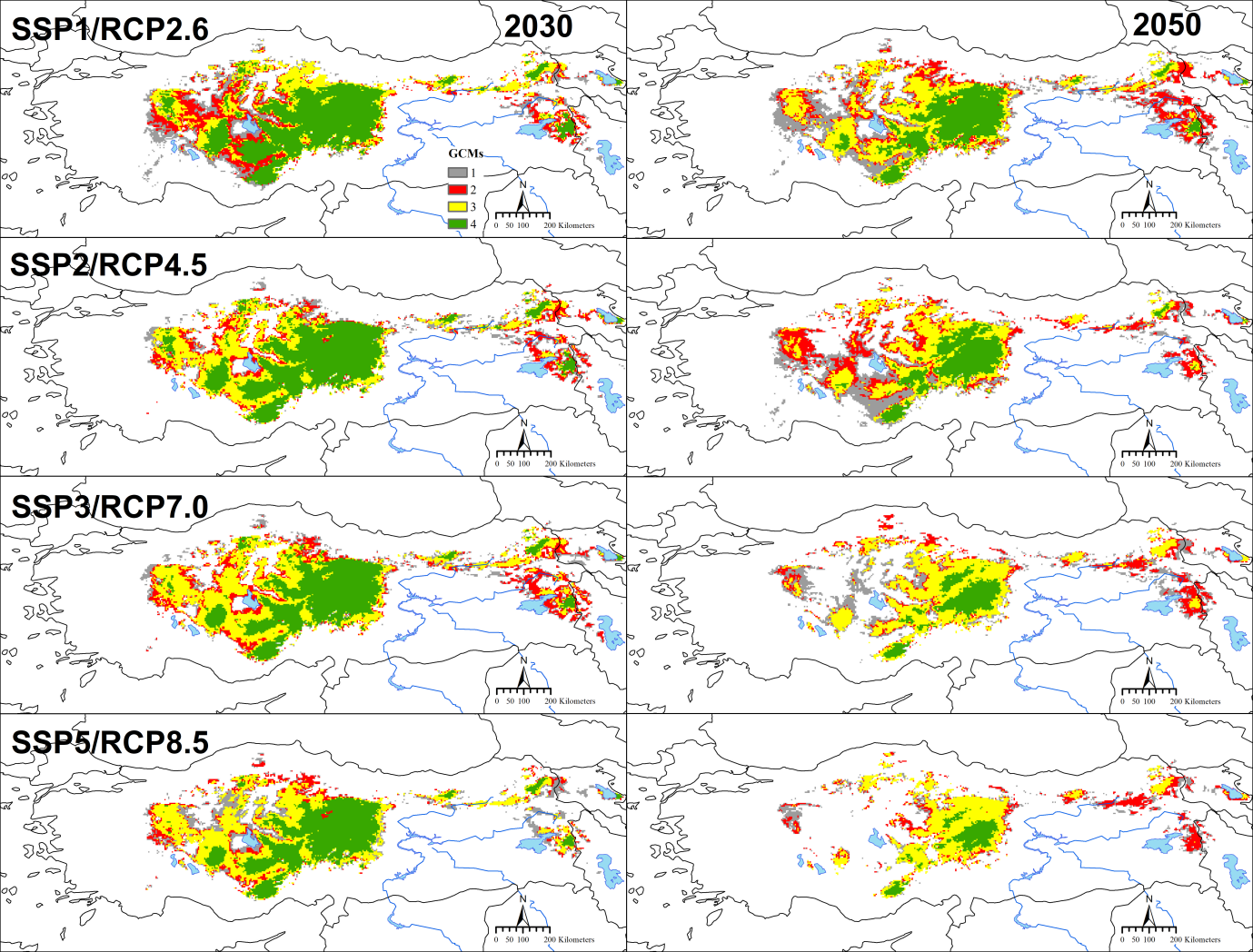

Supplemental Figure 10. Agreement among global climate models (GCMs, BCC-CSM2-MR, CNRM-CM6-1, CanESM5, and MIROC6) in suitable bioclimatic areas under future conditions (i.e. for each scenario, SSP1/RCP2.6, SSP2/RCP4.5, SSP3/RCP7.0, and SSP5/RCP8.5, for each time period, 2030 and 2050)

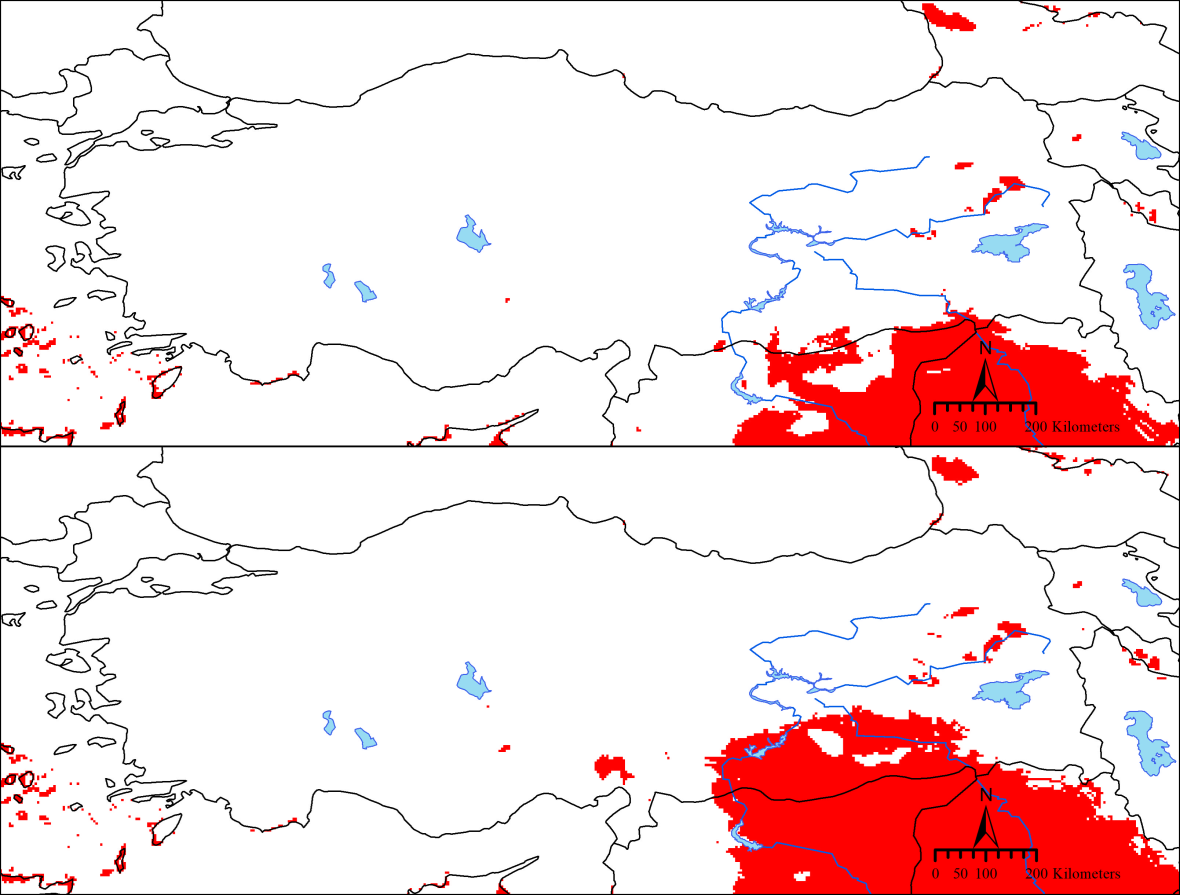

**Supplemental Figure 11.** The results of multivariate environmental similarity surface (MESS) analysis. Non-analog (novel) conditions (for all global climate models and scenarios together, BCC-CSM2-MR, CNRM-CM6-1, CanESM5, and MIROC6 and SSP1/RCP2.6, SSP2/RCP4.5, SSP3/RCP7.0, and SSP5/RCP8.5, for each time period, 2030 and 2050) are shown in red. Above: 2030, below: 2050.

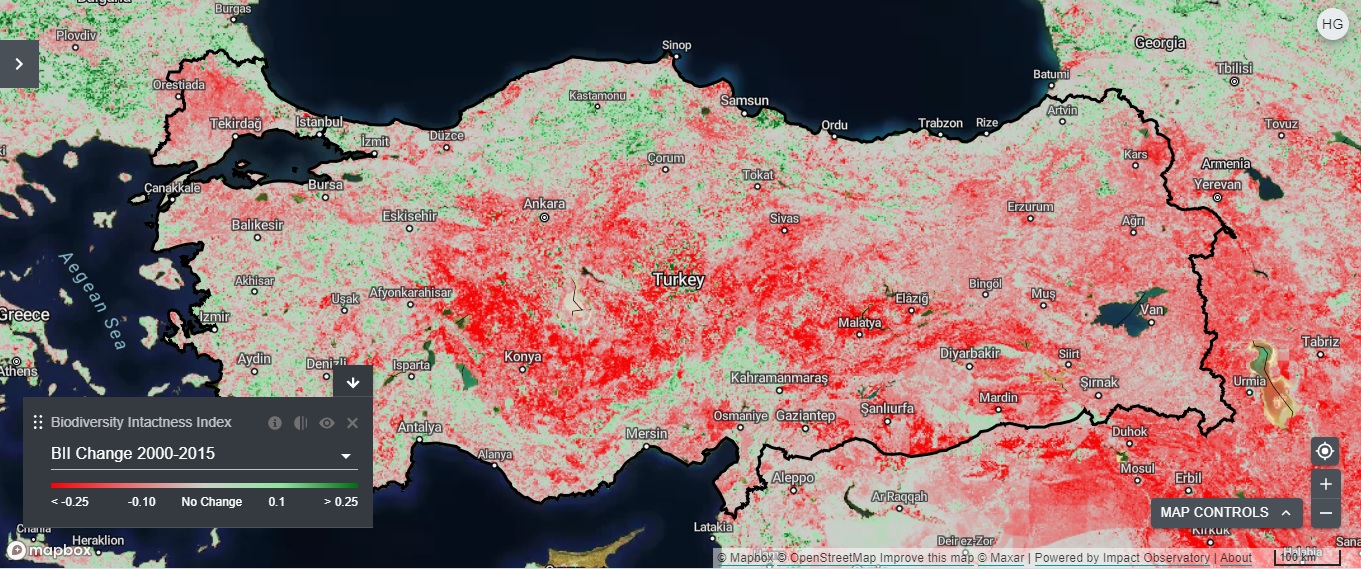

**Supplemental Figure 12.** Biodiversity Intactness Index Change 2000-2015 for Turkey (from UN Biodiversity Lab). A reduction in Biodiversity Intactness Index is shown in red.

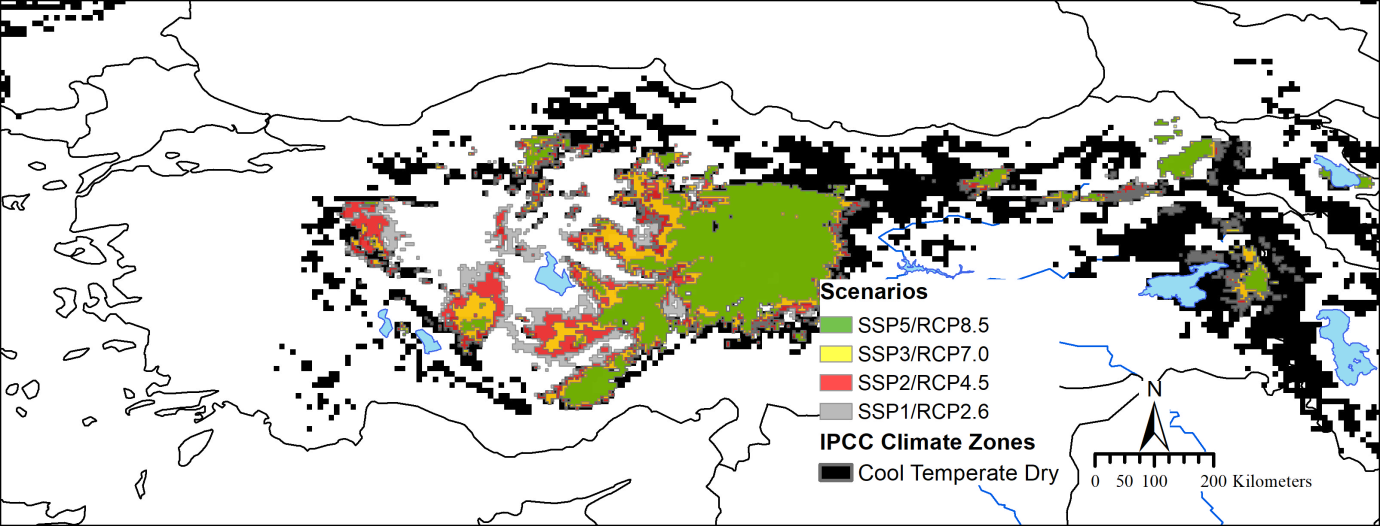

**Supplemental Figure 13.** *In-situ* climate change refugia (i.e. areas predicted suitable both in the present and future, across 2030 and 2050 for each scenario, SSP1/RCP2.6, SSP2/RCP4.5, SSP3/RCP7.0, and SSP5/RCP8.5) for Anatolian ground squirrels (*Spermophilus xanthoprymnus*), with the ‘cool temperate dry zone’ of IPCC Climate Zones (from Joint Research Centre - European Soil Data Centre, ESDAC).

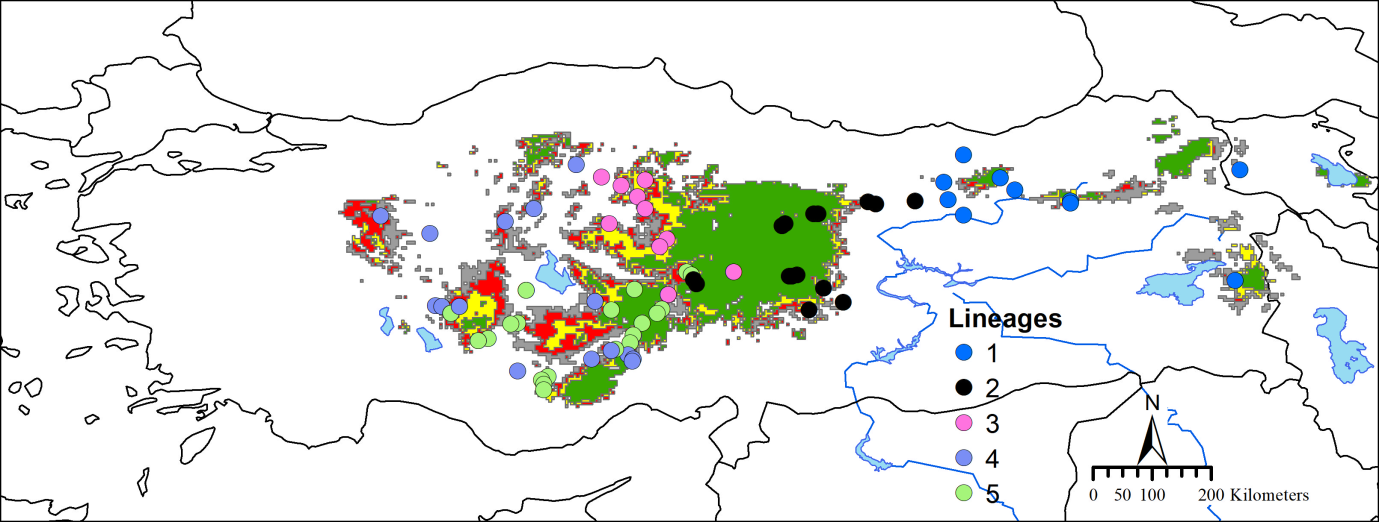

**Supplemental Figure 14.** The geographic distribution of five cytochrome *b* (cyt *b*) mitochondrial (mt)DNA lineages, with *in-situ* climate change refugia (i.e. areas predicted suitable both in the present and future, across 2030 and 2050 for each scenario, SSP1/RCP2.6, SSP2/RCP4.5, SSP3/RCP7.0, and SSP5/RCP8.5), for Anatolian ground squirrels (*Spermophilus xanthoprymnus*). For *in-situ* climate change refugia, see Supplemental Figure 13.

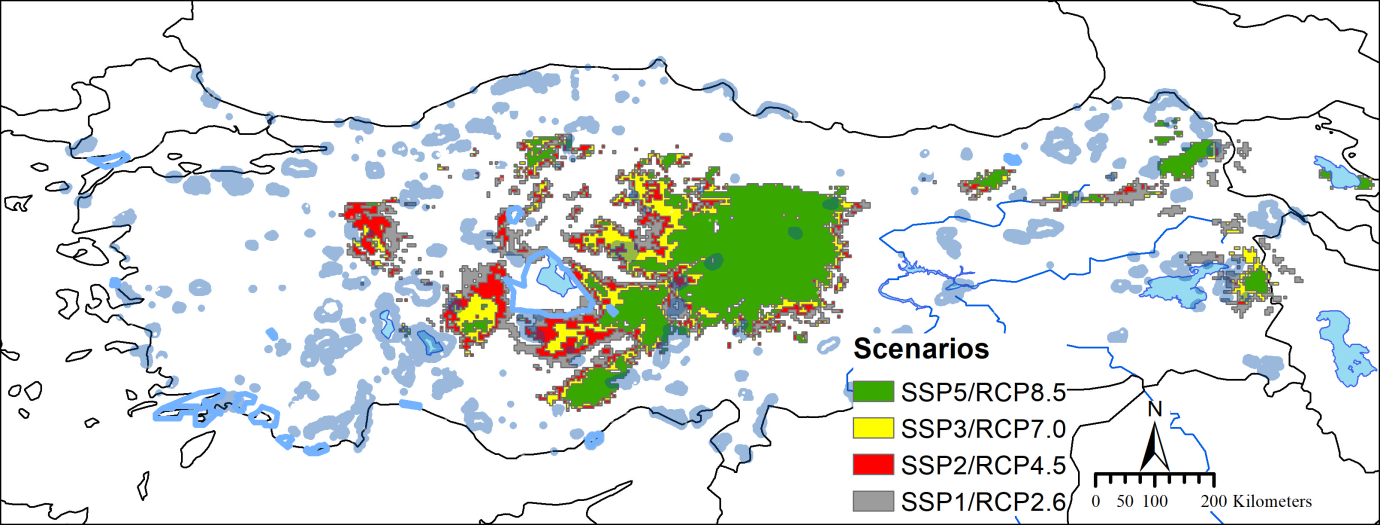

**Supplemental Figure 15.** *In-situ* climate change refugia (i.e. areas predicted suitable both in the present and future, across 2030 and 2050 for each scenario, SSP1/RCP2.6, SSP2/RCP4.5, SSP3/RCP7.0, and SSP5/RCP8.5) for Anatolian ground squirrels (*Spermophilus xanthoprymnus*), with the protected areas (pale blue lines) in Turkey (from Doğa Koruma ve Milli parklar, <https://www.arcgis.com/apps/View/index.html?appid=5f3978146c4643438ab446620e275269> and <http://www.kursatozcan.com/korunan_alanlar/>). represent the
